# Cell-type specific outcome representation in primary motor cortex

**DOI:** 10.1101/2020.03.03.971077

**Authors:** Maria Lavzin, Shahar Levy, Hadas Benisty, Uri Dubin, Zohar Brosh, Fadi Aeed, Brett D. Mensh, Yitzhak Schiller, Ron Meir, Omri Barak, Ronen Talmon, Adam W. Hantman, Jackie Schiller

## Abstract

Adaptive movements are critical to animal survival. To guide future actions, the brain monitors different outcomes, including achievement of movement and appetitive goals. The nature of outcome signals and their neuronal and network realization in motor cortex (M1), which commands the performance of skilled movements, is largely unknown. Using a dexterity task, calcium imaging, optogenetic perturbations, and behavioral manipulations, we studied outcome signals in murine M1. We find two populations of layer 2-3 neurons, “success”- and “failure” related neurons that develop with training and report end-result of trials. In these neurons, prolonged responses were recorded after success or failure trials, independent of reward and kinematics. In contrast, the initial state of layer-5 pyramidal tract neurons contains a memory trace of the previous trial’s outcome. Inter-trial cortical activity was needed to learn new task requirements. These M1 reflective layer-specific performance outcome signals, can support reinforcement motor learning of skilled behavior.

## Introduction

Moving our bodies coordinately in the world is critical for survival. Effective movements require constant adjustment due to changes in the world or the body. To guide movements, the central nervous system must monitor consequences of actions, that can in turn be used to correct ongoing movements and/or inform future movement plans (Keller and Mrsic-Flogel, 2018; Kording and Wolpert, 2006; Tseng et al., 2007; Wolpert et al., 2011).

Action outcomes can be evaluated on two major levels. On one level, outcomes can be evaluated teleologically (i.e., was the ultimate purpose of movement achieved?). Comparisons at this level typically use reinforcement signals such as reward, reward prediction errors, or success and failure outcome signals. On the other level, outcomes can be evaluated with reference to the performance of the physical means of obtaining the goal (e.g., was the planned movement achieved?). Here, movement predictions are compared to sensory consequences of the movement. At this level, feedback error signals serve to indicate sensory prediction errors (Shmuelof and Krakauer, 2011; Wolpert et al., 2011).

Neuronal activity which convey reward, reward prediction errors, and performance error signals are ubiquitous in the brain and have been described in various brain regions including midbrain, cerebellum, and cortical regions (Amador et al., 2000; Amiez et al., 2006; Chen et al., 2017; Engelhard et al., 2019; Heffley et al., 2018; Heindorf et al., 2018; Isomura et al., 2013; Keller et al., 2012; Kostadinov et al., 2019; Krigolson and Holroyd, 2007; Laubach et al., 2000; Mathis et al., 2017; Matsumoto et al., 2007; Roesch and Olson, 2004, 2005; Sajad et al., 2019; Schall et al., 2002; Schultz, 2000; Shadmehr and Krakauer, 2008; Stuphorn et al., 2000; Teichert et al., 2014; Wallis and Kennerley, 2010; Watabe-Uchida et al., 2017; Wickens et al., 2003). Such signals were shown to serve in error-based and reinforcement learning behavioral paradigms (Glascher et al., 2010; Mathis et al., 2017; Shadmehr and Krakauer, 2008; Wolpert et al., 2011) and such combined signals can enhance adaptation (Galea et al., 2015; Nikooyan and Ahmed, 2015).

Representation of motor errors or performance outcome signals in motor cortex (M1) did not receive much attention in the literature until recently. This is despite the fact that M1 is one of the major brain regions responsible for planning and execution of motor commands (Ebbesen and Brecht, 2017; Georgopoulos and Carpenter, 2015; Lemon, 2008; Whishaw et al., 1986), undergoes plasticity at multiple levels, and has been shown to be crucial for motor skill learning (Cichon and Gan, 2015; Fu et al., 2012; Hayashi-Takagi et al., 2015; Li et al., 2017; Makino et al., 2016; Masamizu et al., 2014; Papale and Hooks, 2018; Peters et al., 2014; Peters et al., 2017). Since M1 receives inputs from dopaminergic neurons, sensory streams, and the cerebellum, M1 is expected to have access to both reinforcement and error-based negative outcome signals (Hooks et al., 2011; Hooks et al., 2013; Hosp et al., 2011; Kuramoto et al., 2009; Mao et al., 2011; Molina-Luna et al., 2009; Petrof et al., 2015; Shepherd, 2013; Tsubo et al., 2013).

Three recent studies in primate limb motor cortex demonstrated the involvement of motor cortex in the generation of error signals (Even-Chen et al., 2017; Inoue et al., 2016; Ramkumar et al., 2016). However, the signals reported by the three studies are diverse. Specifically, Even-Chen et al. reported outcome error signals to failure trials during a brain-machine interface based cursor control task. Ramkumar et al. found signals related to lack of reward and not to measures of task outcome during a reaching task to a noisy target. Finally, Inoue et al. reported directionally-tuned visual error signals that were critical for adaptation in a reaching task. Notably, all three studies reported only negative error signals. The presence of only negative error signals in the limb motor cortex is surprising, in light of the importance of positive outcome signals to reinforcement and error based motor learning and the direct dopaminergic innervation to M1 (Galea et al., 2015; Hosp et al., 2011; Molina-Luna et al., 2009; Wolpert et al., 2011). Interestingly, in Purkinje neurons, complex spikes were reported to occur on correct movements and unexpected rewards but not following motor errors during a sensory-motor forelimb task (Heffley et al., 2018), consistent with a reinforcement learning rule. Similar positive signals have not yet been reported in M1 during forelimb behaviors.

While outcome and error signals are beginning to be identified in M1, their nature and circuit realization remains unclear, in part due to the limited number of identified neurons recorded from in the primate. Here by leveraging the tools available in the mouse, we attempt to close this gap and identify movement outcome signals and their realization in M1 networks for a reaching and grasping task. Specifically, we use a head-fixed version of the forelimb prehensile task, consisting of reach-grab-eat in mice (Guo et al., 2015). This is a complex dexterous task, which was shown to be M1-dependent (Guo et al., 2015), and as such facilitates in depth interrogation of the outcome representations in M1. We concentrated on two major excitatory neuronal populations in M1, layer 2-3 pyramidal neurons and layer 5 pyramidal tract (PT) neurons (Anderson et al., 2010; Hooks et al., 2011; Weiler et al., 2008). Layer 2-3 neurons constitute a large cell layer receiving convergent information from cortical, sub-cortical and brainstem regions (Harris and Shepherd, 2015; Shepherd, 2013). On the other hand, PT neurons serve as the main source of motor output to brainstem and spinal cord motor centers (Lemon, 2008). Key open questions include, what type of signals are being generated, are they monitoring reward or the movement itself, can motor cortex generate positive and negative outcome signals, what are the cell type specific and network organization of these signals in M1, and are they useful for motor control and learning during a dexterous task.

Here we show for the first time reward-independent positive and negative outcome signals in layer 2-3 of M1. We find two different populations of layer 2-3 neurons that selectively report either successful or failed behavior attempts. These success and failure outcome signals, report a global assessment of motor performance and not specific kinematic parameters nor reward. Furthermore, outcome impacts PT network initial state activity of the next trial, and post-movement M1 activity is necessary for adaptation to changes in task requirements.

## Results

### Cell-type specific, outcome-related activity in M1 cortex

We imaged layer 2-3 and layer 5 PT M1 neurons (Figure 1) during a prehension task, where head-fixed mice reached for a food pellet, grabbed it, and then delivered it into their mouths. Success was defined when mice grabbed and delivered the food pellet for consumption. This behavior was done without visual information and was cued by an auditory tone with an approximate 30 second inter-trial interval (Figure 1A), as previously described (Guo et al., 2015; Osborne and Dudman, 2014). We used two high speed video cameras (front and side view) and modified machine learning software (Kabra et al., 2013) to semi-automatically annotate the behavior (Figure 1B). We typically annotated three main behavioral epochs, “lift”, “grab”, and “at mouth”, which signify three important phases of prehension in our head-fixed behavioral task (Guo et al., 2015). In addition, we used DeepLabCut (Mathis et al., 2018) to track hand trajectories (Figure S1). Mice were not over trained and on average, mice in this study succeeded on 58±10% of the trials, thus enabling a dynamical range for studying both positive and negative outcome signaling. An average of 2.1 ± 1.3 (n=6 mice; 11 sessions) grab attempts were required to successfully complete the task (Figure 1C and Figure S1). In failure trials, mice typically missed the food pellet at the grab stage or less frequently dropped the pellet before reaching the mouth. On average 3.9 ± 1.7 (n=6 mice; 11 sessions) failed grab attempts were performed before the mice stopped and returned to perch position (Figure 1C and Figure S1). Failures were rarely due to the lack of attempting the prehensile movement.

**Figure 1.**
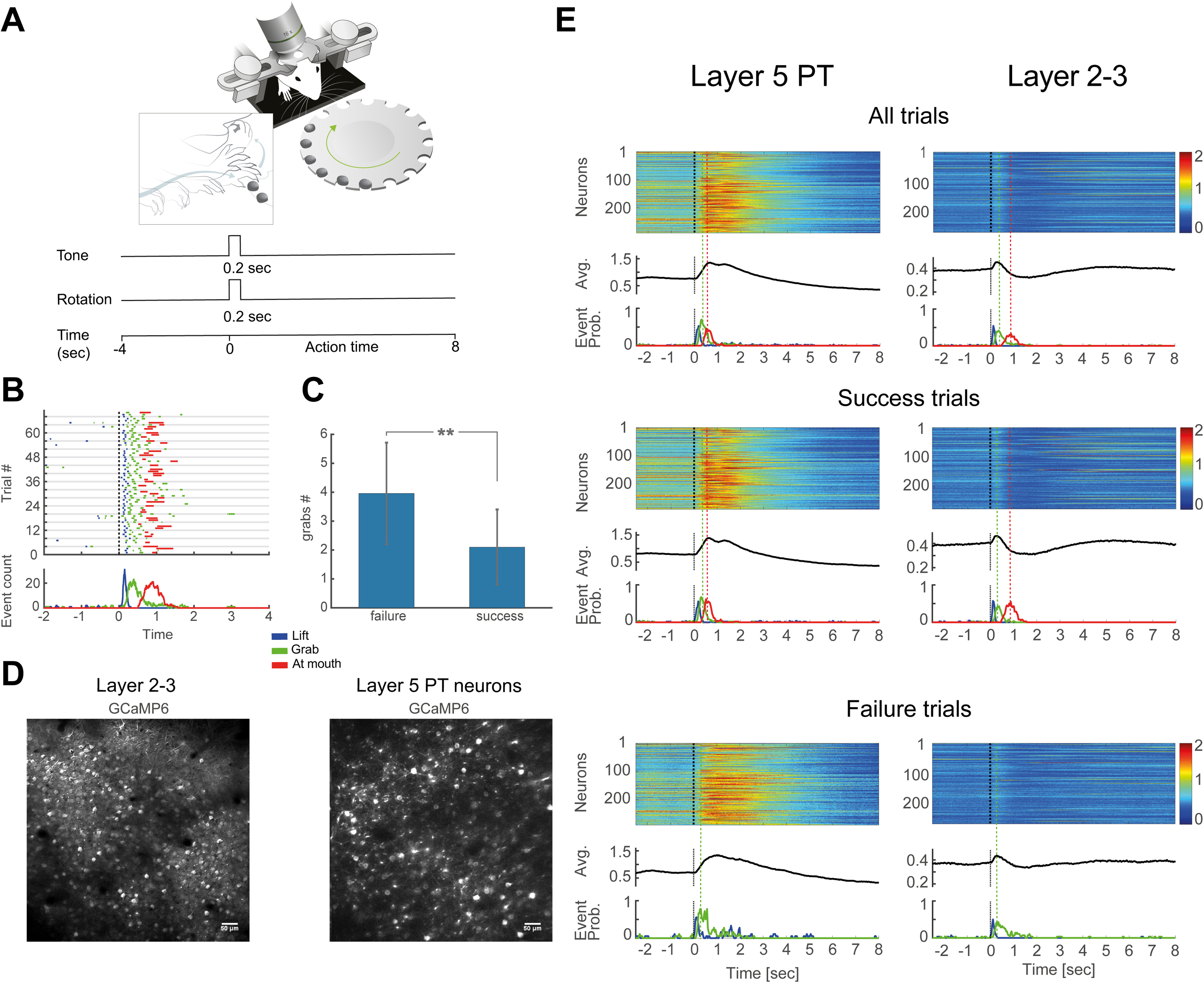
Cell-type specific activity in M1 cortex during prehension. **A**. Behavioral set up. Mice were head fixed and trained to grab a food pellet positioned on a rotating table. Insert, cartoon of the movement. Bottom, trial structure (12 sec). Tone (go signal) after 4 sec, pellet at position at 4.2 sec. **B**. An ethogram annotating the behavior of an expert mouse over consecutive trials (up to 4 sec after tone). Blue, hand lift; green, grab; red, food at mouth. Dashed line represents the tone (black). Bottom panel, peri-event histogram of the behavioral events over time. **C**. Average number of grabs during success and failure trials. 11 sessions over 6 mice (t-test **p<0.01). **D**. Average projections of two-photon imaging of GCaMP6-positive, layers 2-3, M1 pyramidal neurons (230 µm from pia) in a *Slc17a7-IRES-Cre* mouse (left) and layer 5 PT neurons (580 µm from pia) labelled by injection of rAAV2-retro-GCaMP6 virus into the pons of a C57BL6 mouse (right). **E**. Upper panel, averaged (70 trials) calcium transients for all layer 5 PT neurons from a single imaging session during consecutive prehension trials. Color encodes the percent change in fluorescence (ΔF/F). Black trace, grand average over all neurons and trials. Below, event-probability histograms for the behavioral events lift (blue), grab (green) and at mouth (red) aligned to trial onset. Dashed black line denotes the time of the tone. Middle panels, describe success trials and lower describe failure trials. Left column, layer 5 PT neurons, right column, layer 2-3 neurons.

To record activity of pyramidal neurons in M1 cortex, we used chronic, two-photon, calcium imaging of the genetically encoded calcium indicator GCaMP6 (Chen et al., 2017). We used transgenic and viral approaches to record from layer 2-3 pyramidal neurons and layer 5 output PT neurons of M1(Figure 1D-E). Neurons in layer 2-3 were transfected in *Slc17a7-IRES-Cre* transgenic mice (Huang et al., 2013) and PT neurons were specifically retrogradely labeled from the pons in wild type mice using a rAAV2-retro virus expressing GCaMP6 (Tervo et al., 2016) (Figure S2).

If outcome signals exist in M1, they should emerge or be maintained after the behavior and display differences between success and failure trials. On average, layer 5 PT neurons were activated during hand movement and this activity gradually decreased upon the completion of the behavior (Figure 1E). In contrast, the averaged activity of layer 2-3 pyramidal neurons was characterized by a tri-phasic response consisting of an initial, post-tone, brief activation phase; an intermediate, slow inhibition phase; and a late, post-reach, prolonged activation phase (Figure 1E). During the late activation phase, which was coincident or followed behavior completion, a subpopulation of layer 2-3 pyramidal neurons appeared to be specific for the trial outcome (Figure 2A and Figure S3). We identified two main outcome related cell activity patterns: cells that exhibited late activity selectively after successful trials, “success related neurons” and neurons that exhibited late activity after failure trials, “failure related neurons” (Figure 2A). For layer 5 PT neurons, we did not observe comparable late activity (Figure 2B).

**Figure 2.**
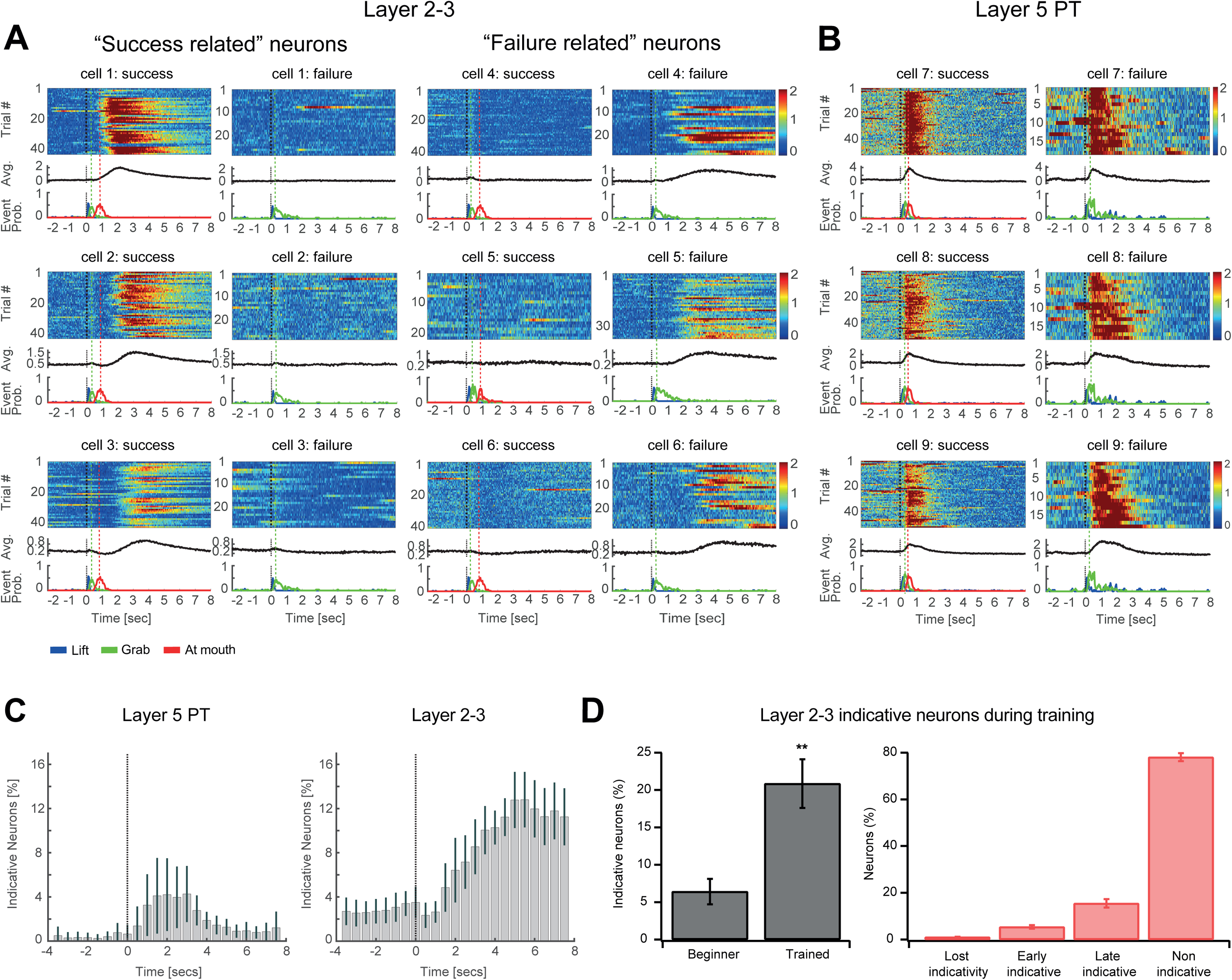
Cell-type specific outcome related activity in M1 cortex during prehension. **A**. Activity of three layer 2-3 “success related” neurons (left panels) and of three layer 2-3 “failure related” neurons (right panels). **B**. Activity of three layer 5 PT neurons. Left column, success trials, right column, failure trials. Trials sorted (ascending) by number of grab attempts. **C**. Indicative neurons were determined using a Support Vector Machines (SVM) classifier that quantified the percent of neurons that reliably reported success or failure trials with 95% confidence at each time bin. Right panel, layer 5 PT (6 mice, 943 neurons) and left panel, layer 2-3 neurons (7 mice, 2293 neurons). **D**. Development of indicative neurons in layer 2-3 during learning. Left panel, average percent of indicative neurons for beginner and trained mice (1640 neurons from 4 mice, t-test p<0.01 compared to the beginner’s phase). Success rates were at 34±4% and 67±7.2% for beginners and trained mice respectively (4 mice, t-test p<0.01 compared to the beginner’s phase). Right panel, percent of neurons (total 1640 neurons) that lost indicativity during training (“lost indicativity”), that were indicative in the beginner phase (“early indicative”), that became indicative in the trained phase (“late indicative”) and neurons that remained non indicative throughout the training (“non-indicative”).

Using a linear support vector machine (SVM) we quantified the percent of layer 2-3 and layer 5 PT indicative neurons that reliably reported success or failure trials (Figure 2C, “indicative neurons” were defined as those that predicted with 95% confidence success or failure trials, see Materials and Method). In layer 5 PT neurons, 6.3 ± 7.4 % were indicative during the initial 3 seconds after the tone, but gradually diminished during the later phase of the trial (Figure 2C; 6 mice). Since the hand movements occurred mostly during the initial 3 seconds after the tone, the calcium activity of some of these indicative neurons could in fact reflect movement differences between successes and failures such as the number of grab attempts (Figure 1C). In contrast, in layer 2-3 neurons, approximately a sixth of the layer 2-3 population were indicative (16.1 ± 5.8 %; 7 mice). Their activity gradually developed during the first 5 seconds after the tone and remained high throughout the duration of the trial (Figure 2C). Therefore, late signals that distinguish outcomes are found in layer 2-3 M1 neurons.

Are outcome neurons innate in the layer 2-3 network or do they develop as the task is learned? To answer this question, we compared indicative neurons at two time points during the training, when mice were beginners with average success rate of 34±4% and later in the training when they almost doubled their success rates (67±7.2%; 4 mice, t-test p<0.01 compared to the beginner’s phase). We found that indicative neurons developed with training, increasing from 6.4±1.7% to 20.9±3.3% of the recorded neuronal population (Figure 2D; 4 mice, 1640 neurons, t-test p<0.01). Only a small minority ∼ 1% lost their indicatively from the early to the later time point of training (Figure 2D). These findings indicate that the majority of outcome related neurons are not innate residents of the layer 2-3 network of M1, rather they develop as part of learning process.

### Layer 5 PT neuronal activity is primarily movement related while layer 2-3 neuronal activity is primarily outcome related

We next used latency histograms to examine the relationship between movement parameters and neuronal activity. We quantified the percent of neurons with a peak response in different latencies relative to the various behavioral events. A large percentage of layer 5 PT neurons showed locking to tone and the different behavioral events such as first and last grab, and hand at mouth (Figure 3A). In contrast, a small number of layer 2-3 neurons showed locking to the tone, lift, and first grab and locking was nearly absent for the last grab (Figure 3B, Figure S3 and Figure S4). The average percent of layer 5 PT and layer 2-3 neurons activated within 0.2 seconds before and 0.5 seconds after the first grab was 67.8±7.8% and 9.9±2.1% and the last grab was 49.3±7.7% and 4.7±0.9% respectively (4 animals for layer 2-3 neurons and 4 animals for PT neurons). Therefore, unlike PT activity, the outcome-related activity of layer 2-3 was not well temporally aligned with motor behaviors.

**Figure 3.**
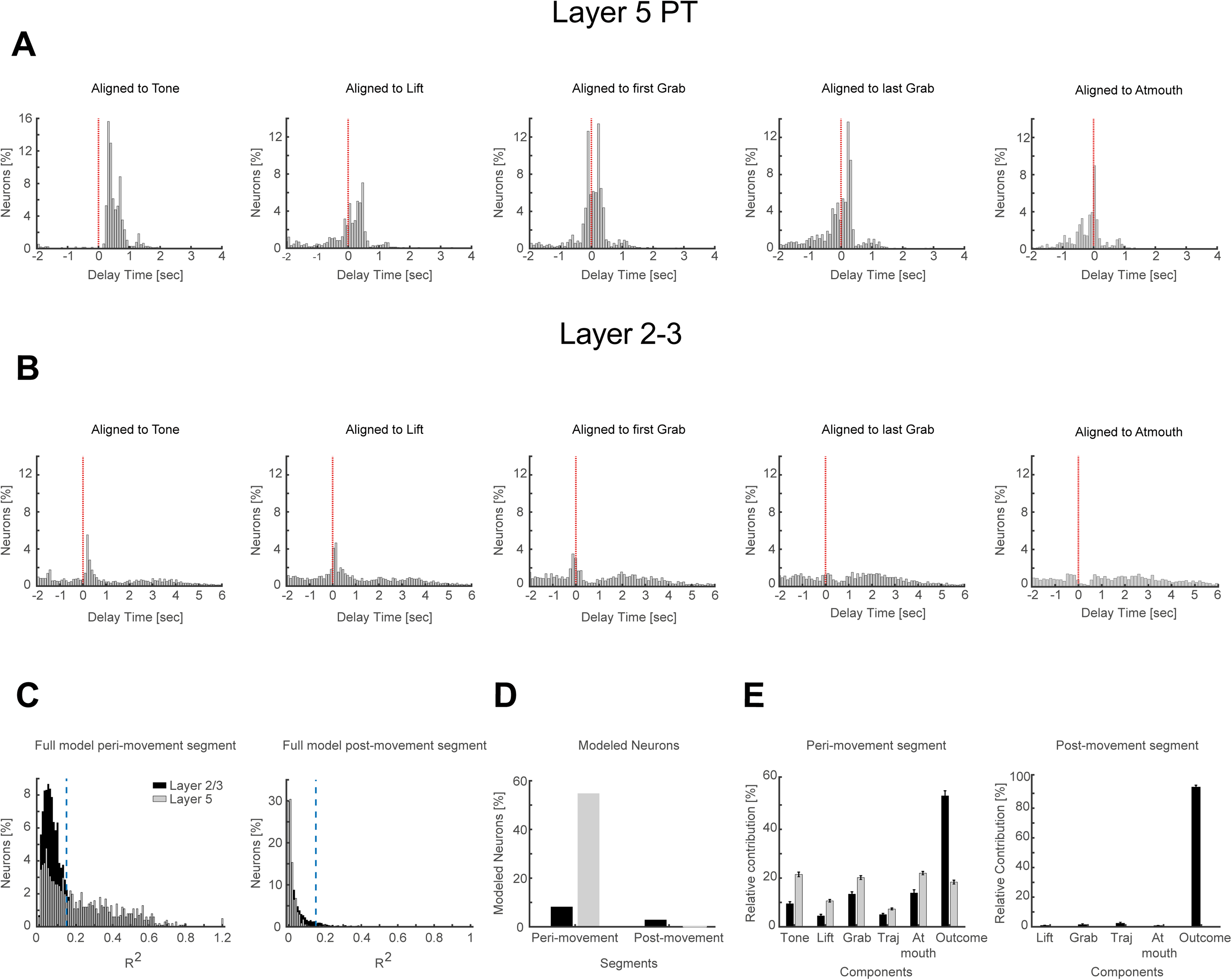
Relationship between movement parameters and layer 2-3 and layer 5 PT activity. **A**. Latency histograms of percent peak layer 5 PT neuronal activity (4 animals; 1045 neurons; 375 trials) aligned to tone, lift, first grab, last grab and at mouth (left to right). **B**. Same as in A, but latency histograms of layer 2-3 neuronal activity (4 animals; 2063 neurons; 1052 trials). **C**. Modeling calcium transients of layer 5 PT and layer 2-3 neurons using a generalized linear model (GLM). Distribution of percent of neurons (1008 neurons for layer 5 PT and 1111 neurons for layer 2-3) as a function of their R^2^ (values averaged over 5-fold cross validation on each time segment; left, for peri-movement time segment (tone -1 to tone +2 sec) right, for post-movement time segment (tone +5 to tone + 8 sec). **D**. Percentage of neurons included in the analysis, i.e. neurons in which at least 15% of the variance was modeled by the GLM. **E**. Bar graph of the relative contribution of modeled neurons (with at least 15% of their variance explained) for each of 6 predictors (tone, lift, grab, at mouth, hand trajectories, success/failure trial status) composing the full model. Left, peri-movement time segment, right, post-movement time segment.

To further quantify the relationships between motor events and neuronal activity in both layers, we used a generalized linear model (GLM) (Engelhard et al., 2019; Ramkumar et al., 2016) to evaluate the relative contributions of the different behavioral components (see Methods) to the observed calcium signals. We modeled the calcium transients of layer 2-3 and layer 5 PT neurons based on 3 types of predictors: time series of 3-D spatial location of the hand (Mathis et al., 2018), time-varying discrete behavioral events (tone, lift, grab, and at mouth) (Kabra et al., 2013), and outcome labeled as constant binary variables of either success or failure (1 or 0 respectively) throughout the duration of the trial (Figure S5). To model the time course of the GCaMP6 calcium signal of single neurons, we convolved the time varying binary events with a set of splines (Figure S5A-B). We modeled the neuronal activity as a linear combination of the predictors during two temporal epochs: peri-movement segment during which mice complete the hand reach attempts (1 second before until 2 seconds after the tone; Figure S5C-D); and post-movement segment (5 to 8 seconds after tone). For the peri-movement time segment, the GLM was more successful in modeling the activity of individual PT neurons than of layer 2-3 neurons. During this first time segment, the full GLM successfully modeled more than 15% of the energy in 51.5% of layer 5 neurons and in only 4.5% of layer 2-3 neurons (Figure 3C-D). Motor related parameters were more prominent in layer 5 PT neurons (explaining ∼75% of the modeled activity), while outcome parameters were more prominent in layer 2-3 neurons (explaining >50 % of the modeled activity) (Figure 3E). In contrast to the peri-movement segment, during the post-movement time segment, the full GLM could not model any of the layer 5 PT neurons while successfully modeling the activity of ∼ 3% of layer 2-3 neurons (Figure 3E). We presume that the difference in the percent of neurons modeled by the GLM compared to the larger number of indicative layer 2-3 neurons, resulted from two fundamental assumptions used by the GLM: linear interactions between the different predictors and uniform responses for trials of any given predicted parameter. For layer 2-3 neurons with post-movement modeled activity, the outcome parameter explained nearly all (∼95%) of the activity (Figure 3E). Taken together our analysis revealed that while a major fraction of the activity of layer 5 PT neurons was related to movement itself, outcome information explains more of the activity in layer 2-3 especially in the post-movement time segment.

### Differential network representation of outcome-related activity in layer 5 PT and layer 2-3 neurons

Next, we investigated how the outcome related activity is represented at the network level. First, we performed principal component analysis (PCA), and embedded the data from all neurons of each trial into a 2D plane defined by the first two principal components obtained by the PCA (Figure 4A-B). Thus, each point in this analysis represents the associated values in the two principal eigenvectors of a single trial. The PCA embedding of the data obtained for layer 2-3 could discriminate between success and failure trials with a very high level of accuracy (Figure 4A; average accuracy 0.95±0.05; 8 mice; 24 sessions). Interestingly, the algorithm picked up unusual behaviors, for example, when the mouse groomed during the trial (Figure 4A, orange dots). The second dimensionality reduction analysis method performed an unsupervised nonlinear embedding of the data based on manifold learning (diffusion maps) (Lafon et al., 2006) in order to create a hierarchical tree-like clustering (see Methods). When we applied this analysis to the data of layer 2-3 neurons, failure and success trials separated at the root of the tree, implying that the initial separation of the tree was based on the trial outcome (Figure 4A; Average accuracy 0.95±0.06; 8 mice 24 sessions). Taken together, these two analysis methods revealed that motor outcome is represented in the top activity components of the M1 layer 2-3 network.

**Figure 4.**
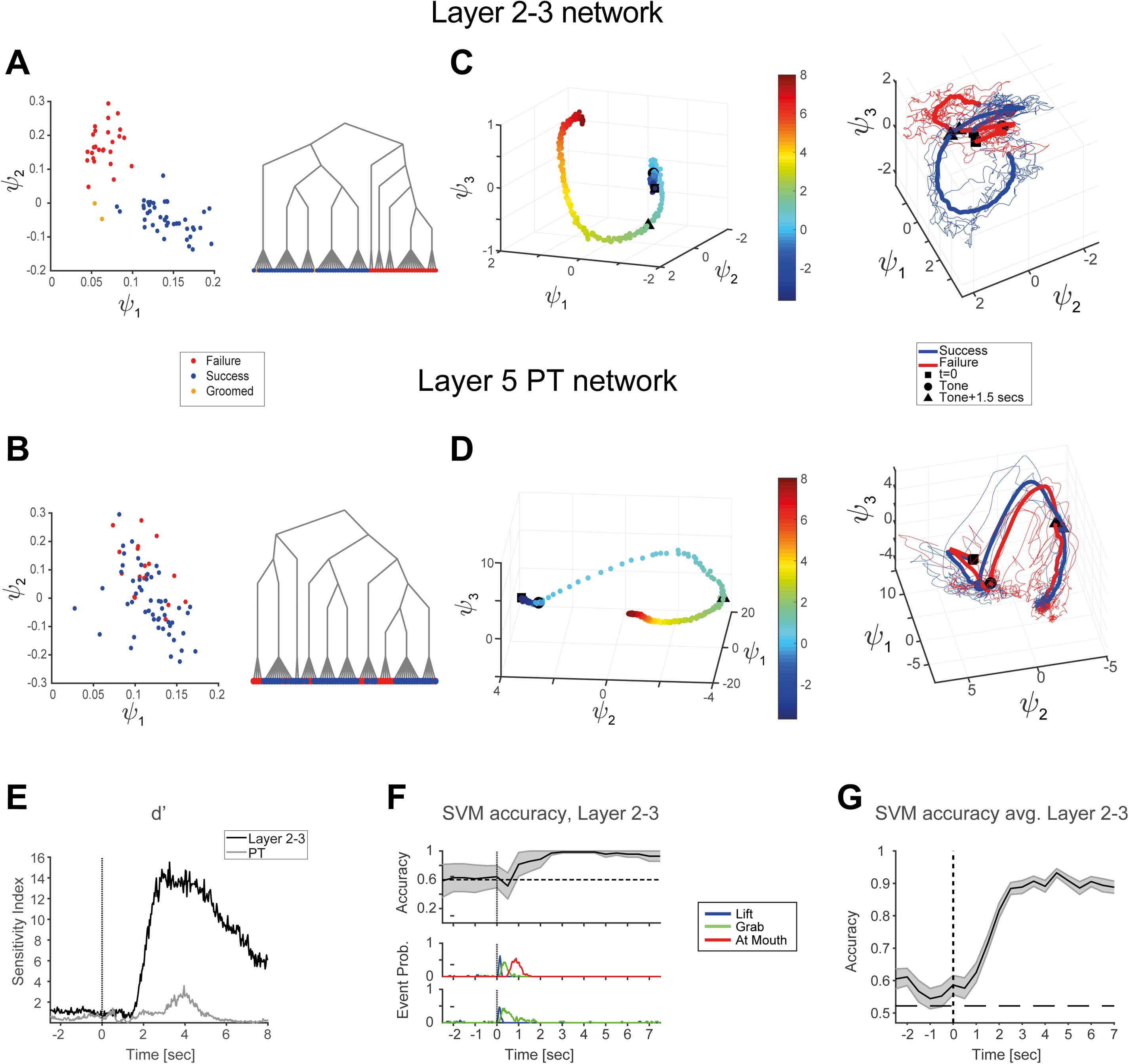
Outcome related activity of layer 2-3 and PT networks during prehension. **A**. PCA embedding of layer 2-3 neurons from one session (70 trials). The 3D matrix of neuronal activity over time and trials was reduced into a 2D matrix of all neurons at all times for all trials using PCA. Trials are represented in first two PC space. (Red, failure; blue, success; orange, trials with grooming). Accuracy of separation was 0.97±0.06. Right, diffusion map embedding of same data points. **B**. PCA embedding of layer 5 PT neurons from one session. Accuracy of separation was 0.74±0.17. Right, diffusion map embedding of same data points (color code same as in A). **C**. Left panel, averaged temporal evolution of layer 2-3 network activity projected over the first three PC axes. Each PC is a linear combination of the neuron’s dimensional space (274 neurons in this example; same data as in A; color depicts time). The starting point (square), tone (circle) and 1.5 seconds post tone (triangle) are marked along the trajectory. Right panel, average temporal evolution of the network activity separated to success (blue) and failure (red) trials for layer 2-3 neurons (same experiment as in A). Examples of single trajectories each representing a single trial are overlaid. Average trajectory in bold. **D**. Left panel, average temporal evolution for PT neuronal activity (same data as in B). Right panel, average temporal evolution of the PT network activity separated to success (blue) and failure (red) trials (same experiment as in B). Examples of single trajectories, each representing a single trial are overlaid. Average trajectory in bold. **E**. Sensitivity index (d’) calculated to quantify the separation between the two trajectories (success and failure) in the different time windows, shown for layer 2-3 (black line) and for PT neurons (grey line). **F**. Top, SVM accuracy of success/failure trial separation for layer 2-3 activity (same experiment as in A, C) calculated in 1 sec time bins, with 0.5 sec overlap, using 10-fold cross validation (± SD shown in grey). Bottom, event-probability histograms for the behavioral events lift (blue), grab (green) and at mouth (red). Dashed black line denotes the time of the tone. **G**. Average SVM accuracy for success/failure classification, calculated for layer 2-3 neuronal activity in 8 mice, 25 sessions.

Similar PCA embedding analysis performed for layer 5 PT neurons also showed classification of the success and failure trails albeit with lower accuracy (Figure 4B; Average accuracy 0.74±0.11, 8 mice, 25 sessions). However, given the correlation in the latency histograms and the GLM analysis (Figure 3), this classification may reflect at least in part the behavioral differences in success vs. failure trials which exhibit more grab attempts (Figure 1C and Figure S1). Diffusion maps embedding of layer 5 PT neuronal activity showed that in contrast to layer 2-3 neurons, outcome information is not represented in the primary division of the tree, rather at lower branching levels indicating that outcome is not the primary information content of layer 5 PT neurons (Figure 4B; Average accuracy 0.77±0.16, 8 mice, 25 sessions).

Since M1 is likely a dynamical system (Churchland et al., 2012; Cunningham and Yu, 2014), we examined the temporal evolution of the compressed layer 2-3 and PT network activity during the hand reach trials (see Methods for more details). The activity was plotted with the first three PC values represented by the three axes of the plot, thus visualizing the 3D dynamical evolution of the network during the time course of the trial (Figure 4C-D). For layer 2-3, these three PC value represented 66±2.8 % of the total activity energy, and average of 46.6±6.7 eigenvalues were required to account for 95% of the total activity energy (n=15 sessions). For layer 5 PT neurons, the initial three PC value represented 88.5±2.7 % of the total activity energy, and average of 15.9±4.2 eigenvalues were required to account for 95% of the total activity energy (n=13 sessions). Abrupt changes in the layer 2-3 and layer 5 PT neural trajectory in state space were aligned to behavioral events (Figure 4C-D), for example tone (4 sec, circle) and reach completion (∼ 5.5 sec, triangle). When we plotted the average trajectories of failure and success trials, we observed that these trajectories for both layer 2-3 and layer 5 PT neurons started in close proximity in state space and developed in parallel up to the tone. However, for layer 2-3, the trajectories of successful and failure trials diverged after the tone (Figure 4C). In contrast, layer 5 PT trajectories developed in close parallel paths throughout success and failure trials (Figure 4D). To quantify the degree of separation between the trajectories obtained for success and failure trials in layer 2-3 and layer 5 PT neurons, we used a sensitivity index (d’) calculated for the first three PC components. In layer 2-3 neurons the d’ showed a significant deflection in the separation of success and failure trials late in the trial after most hand reach movements terminated (peaking ∼ 3 sec after the tone; Figure 4E). The separation of the PC trajectories was consistent across all animals tested (8 mice, 25 sessions; average peak of the d’ was 16.83±11.84). In contrast, for layer 5 PT neurons, the degree of separability as measured by d’ was smaller compared to layer 2-3 pyramidal neurons (Figure 4E; n=8 mice, 25 sessions; average d’ was 4.68±2.4). The small separation observed for layer 5 PT neuron trajectories, suggest the layer 5 PT network does represent outcome information, but at a much smaller magnitude and for a shorter period of time than the layer 2-3 network. These findings are consistent with our SVM indicative neuron and GLM analysis (Figure 2C and 3E). To test the separability of layer 2-3 network activity for success and failure trials based on the entire network activity, rather than based on the first three PCs, we used an SVM approach. This analysis also showed that layer 2-3 could discriminate the outcome of the previous action, with high accuracy (average accuracy for layer 2-3 neurons was 0.96 ± 0.03 for 8 mice, 25 sessions) after ∼ 1.5 sec following the auditory cue when most hand reach related movements ended (Figure 4F-G). Therefore, the layer 2-3 network best monitors outcome of recent actions and the distinguishing signals are most significant after the completion of behavior.

### Outcome-related activity in layer 2-3 neurons of M1 depends on motor performance and not on food consumption/reward

We next performed behavioral modifications to decouple hand reach movements and food consumption/reward. To test the necessity of hand movement, the behavioral task was modified such that instead of performing a hand reach for pellet, the mice licked (“tongue reach”) for the food pellet (Figure 5; movie 1 and 2). In these experiments, hand reach mice were switched to the tongue reach paradigm (Figure 5A). The results of these experiments showed that once food was accessed via the tongue the overall average activity was significantly attenuated (Figure S6A-C). Importantly, almost all outcome success- or failure-related neuronal activity disappeared (Figure 5B-C) and the percentage of indicative neurons markedly decreased (3.7 ± 3.1% during tongue reach and 14.1 ± 4.3 % during hand reach; 5 mice, 14 sessions; p<0.01; Figure 5D-E). Furthermore, the accuracy of discriminating between success and failure trials decreased to chance levels in the tongue reach task (Figure 5D-E; 4 mice). Thus, food consumption or reward signals are not sufficient to generate the outcome related activity observed in layer 2-3 of M1.

**Figure 5.**
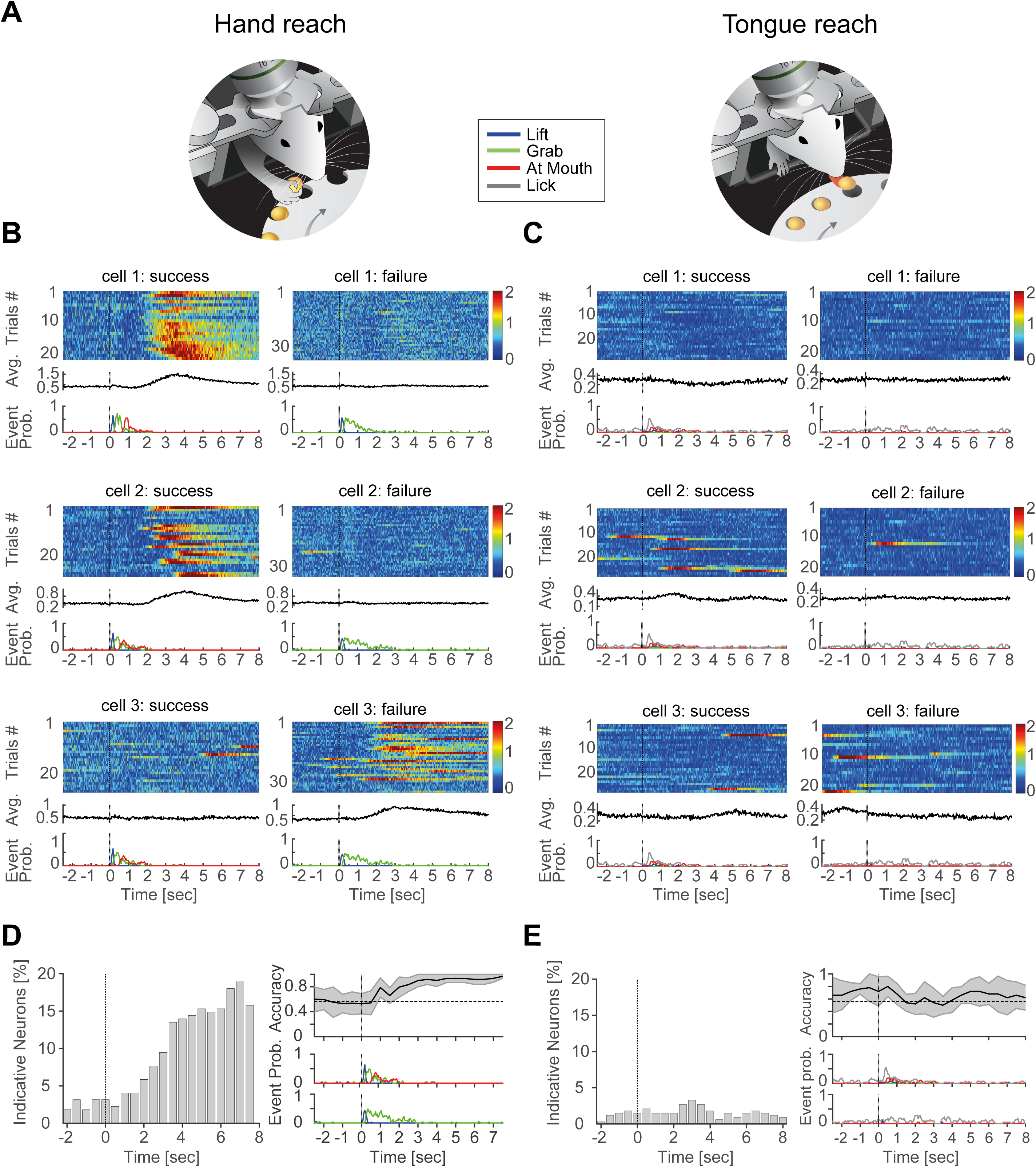
Comparison of neuronal activity during hand and tongue reach experiments. **A**. Mice were trained for hand reach (left) and later trained for tongue reach (right). Trial structure was identical. **B**. Activity of three layer 2-3 neurons during the hand reach behavioral paradigm. Activity during success (left column) and failure (right column) trials during one session. **C**. Same neurons as in B during a tongue reach session. **D**. Percentage of indicative neurons (left), SVM accuracy (right) for hand reach. **E**. Percentage of indicative neurons (left), SVM accuracy (right) for tongue reach.

To test if performance of the behavior was sufficient to produce the outcome signals without reward, we exchanged the food pellets with non-edible, 3-D printed, plastic pellets. To provide a similar sensory experience, the plastic pellets had similar size, shape and weight to the normal pellet. Typically, in these experiments the mice performed the entire hand movement including bringing the plastic pellet to the mouth but never swallowed the plastic pellets (see movie 3). The definition of failure trials did not change in these experiments, however, success trials were defined when the mouse grabbed and brought the plastic pellets to the mouth. In these experiments despite the fact that no reward was actually consumed, we still observed outcome related neurons with a high percentage of indicative neurons (Figure 6A-D; 37.9±13.3%; 5 sessions in 2 mice). Furthermore, at the network level success and failure trials were clearly distinguished by the layer 2-3 network activity. The 2D PCA embedding could reliably discriminate between success and failure trials with accuracy of 0.94±0.06 for plastic pellets (5 sessions in 2 mice; Figure 6F) compared to 0.95±0.05 for normal pellets (4 sessions in the same 2 mice; Figure 6E) and the SVM classified the outcome related activity with high accuracy for the normal and plastic pellets (0.96±0.01 for plastic pellets and 0.92±0.04 for normal pellets, 95% confidence; 5 sessions in 2 mice and 4 sessions in 2 mice respectively). Similar to the normal pellets, the average layer 2-3 neural trajectories completely separated and evolved in different directions during failure and success trials (Figure 6F; d’ was 19.6±4.7 for plastic pellets and 17±1.4 for normal pellets, 5 sessions in 2 mice and 4 sessions in 2 mice respectively). Taken together, the tongue reach and plastic pellet experiments indicate that outcome-related neuronal activity is not dependent on food consumption, but rather seems to reflect an evaluation of the motor performance itself.

**Figure 6.**
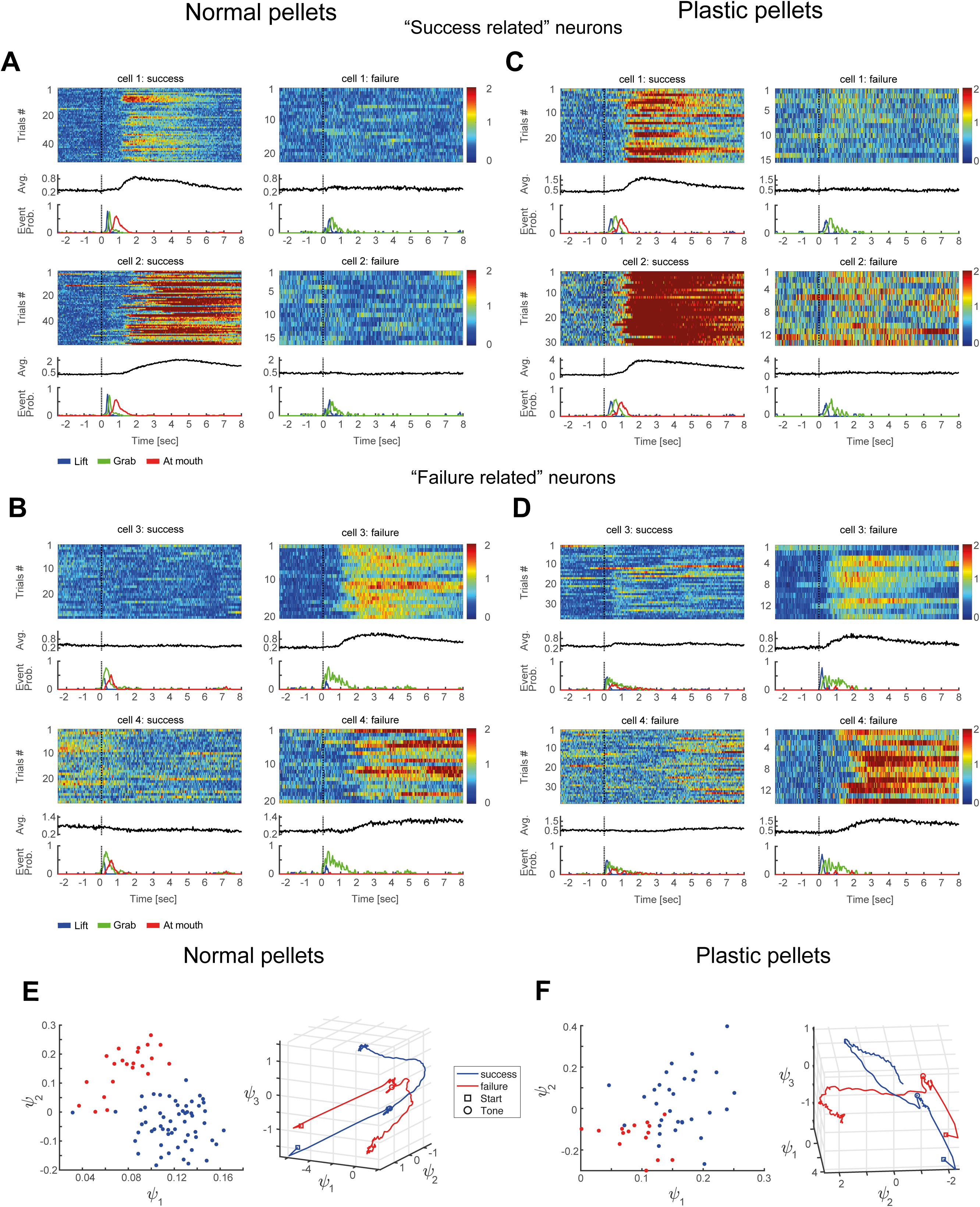
Outcome related signals in layer 2-3 neurons in the absence of appetitive reward. **A**. Activity of two layer 2-3 success related neurons during hand reach trials with normal pellets. **B**. Activity of two layer 2-3 failure related neurons during hand reach trials with normal pellets. **C**. Activity of two layer 2-3 success related neurons during hand reach trials with plastic pellets. **D**. Activity of two layer 2-3 failure related neurons during hand reach trials with plastic pellets. Example neurons are the same for plastic and normal pellets. **E**. Left panel, 2D PCA embedding for layer 2-3 neurons with normal pellets in one example experiment (red, failure; blue, success). Accuracy of separation 0.96±0.07. Right panel, average temporal evolution of the network activity separated to success (blue) and failure (red) trials for layer 2-3 neurons with normal pellets. **F**. Left panel, 2D PCA same as in E, for plastic pellets. Accuracy of separation 0.94±0.09. Right panel, average temporal evolution same as in E, for plastic pellets. Same animal for normal and plastic pellets.

Motor performance could be evaluated based upon the trajectory of the arm to the pellet or whether a food pellet was successfully grabbed. To test if touching the food pellet was required for success- and failure-related signals, we omitted the pellet on a fraction of trials (24.5 ± 2.7 % of the trials, n=3 mice; 11 sessions). Omission trials nearly exclusively led to activity in failure-related neurons (Figure S7A). The PCA embedding of the data obtained for layer 2-3 could not discriminate between omissions and failure trials (Figure S7B; 0.61±0.12, chance 0.63±0.06; 95% confidence). Furthermore, the average layer 2-3 neural trajectories of failure and omission trials were close together and the discrimination accuracy using a SVM was at chance level (Figure S7C-D; 0.6±0.09; chance 0.62±0.06). Taken together, the three behavioral manipulation experiments we performed show that success- and failure-related signals must reflect the experience or absence, respectively, of a post-grab, non-reward, sensory event.

### Outcome impact the initial state of layer 5 PT network

If outcome signals are to be useful, there should be some memory of these signals when future behavior is being produced. If this is indeed the case, future neuronal activity could change according to the performance of the preceding trial. To investigate this possibility, we compared the average trajectories of the layer 2-3 and layer 5 PT network activity during hand reach trials according to both the present and previous outcome of the trials (Figure 7A-D). Specifically, we divided the trials into 4 subgroups: success following success, success following failure, failure following success and failure following failure. The initial state of the layer 2-3 network showed no dependency on the outcome of the preceding trial (Figure 7A). The accuracy of determining the outcome of the previous trials based on the ongoing activity of the layer 2-3 network was at the chance level throughout the course of the trial (Figure 7B; average accuracy 0.65±0.05, chance 0.63±0.07; calculated 2 seconds before the tone, 14 sessions in 4 animals that passed the inclusion criteria of at least 30 trials in each outcome group per session). The confusion matrix showed the inability of the SVM classifier to identify previous labels of success and failure trials. In contrast, the initial state of the layer 5 PT network was affected by the outcome of previous trials (Figure 7C). The initial pre-tone state of success-success and success-failure trials were in close proximity to each other (blue and purple trajectories), as were the pre-tone state of failure-success and failure-failure trials (cyan and red trajectories). As the hand reach movement progressed the trajectories of the pairs in these two groups parted and formed two new groups, success-success and failure-success pairing (blue and cyan trajectories) and success-failure and failure-failure pairing (purple and red trajectories) based on the present movement outcome (Figure 7C and movie 4). The accuracy of determining the outcome of the previous trial based on the ongoing activity of the PT network was significantly higher than chance (Figure 7D; Average accuracy 0.71±0.09, chance 0.58±0.07; p<0.01; calculated 2 seconds before the tone, 9 sessions in 4 animals that passed similar inclusion criteria as layer 2-3 sessions). The confusion matrix showed that the SVM classifier had comparable accuracy of correctly identifying both preceding success and failure trials (Figure 7D). Evidence for memory of past trial outcome on the initial activity state of the present trial was not only observed at the network level, but also on single neuron resolution as well (8.5±4.46 % of neurons, 95% confidence, 4 animals in 14 sessions) (Figure 7E). Taken together, our findings indicate that initially outcome is determined by layer 2-3, but by the next trial this information is represented in the PT network.

**Figure 7.**
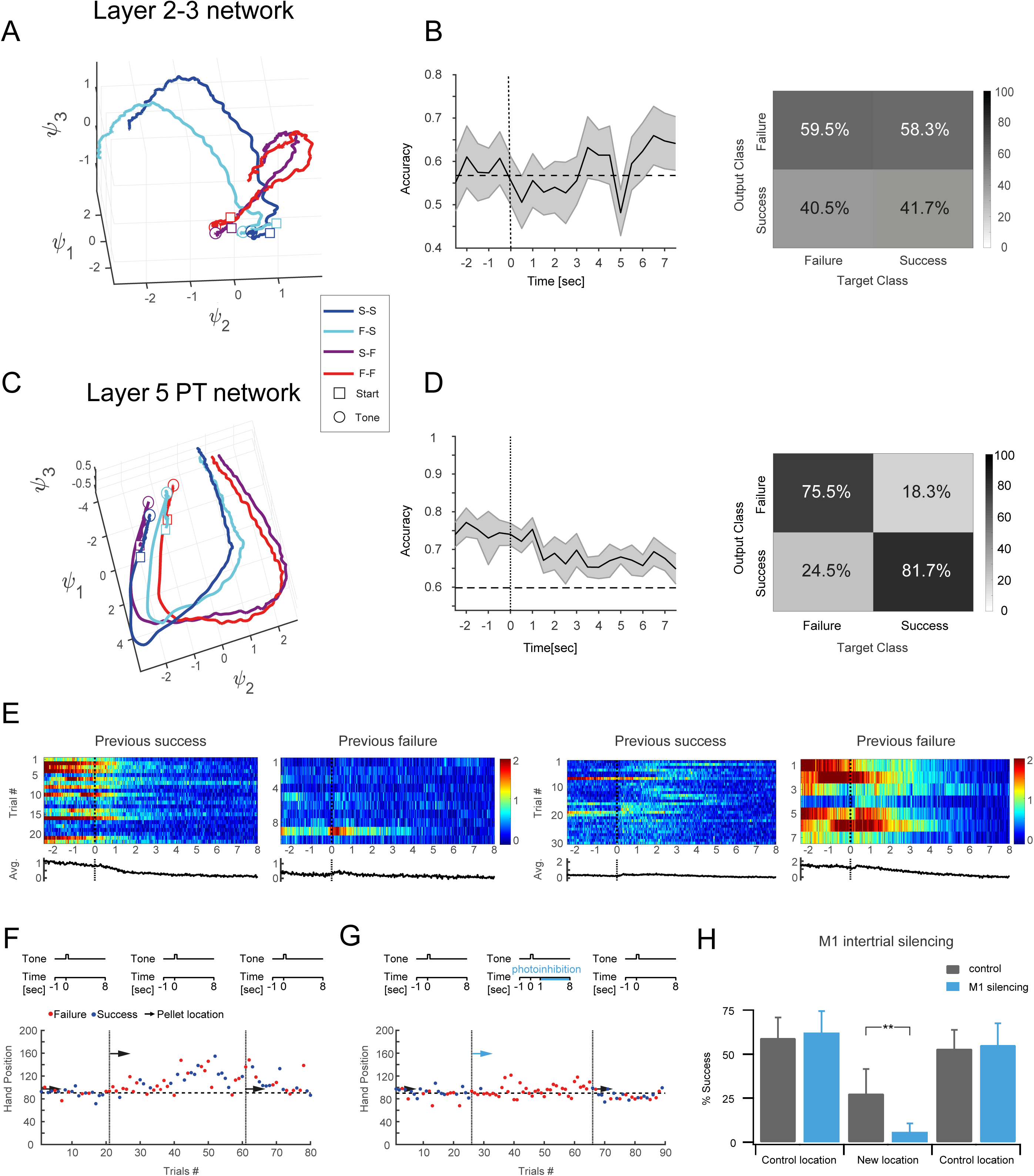
Memory and utility of outcome related signals in M1. **A**. Average temporal trajectories of layer 2-3 neurons were sorted according to previous and present trial outcomes resulting in 4 types of trajectories: previous and present successes (S-S, blue); previous failure and present success, (F-S, cyan); previous success and present failure, (S-F, violet); previous and present failures (F-F, red). Square marker denotes start of trial; circle denotes the time of the tone (4 sec after start). **B**. Left panel, SVM classifier (± SD shown in grey, dashed horizontal line represents chance level) for the data shown in A, did not successfully predict the behavioral outcome based on the activity of the next trial. Right panel, confusion matrix calculated for success and failure trials. **C**. Average temporal trajectories of layer 5 PT neurons, were sorted according to previous and present trial outcomes. Sorting as described in A. **D**. Left panel, SVM classifier for the data shown in C, successfully identified the behavioral outcome based on the activity of the next trial. Right panel, confusion matrix calculated for success and failure trials. **E**. Example of PT neurons sorted according to previous success trials (left) or failure trials (right). **F**. Hand position on successive trials (blue success, red failure) in control pellet location, after movement of the pellet and back to control pellet location (arrow represents pellet location). **G**. Same as in F, except at trials where the pellet was moved to a new location, photoinhibition was activated 1000 ms after the tone (see scheme above). Same animal for F-G. **H**. Summary plot for success rate during control, new pellet location and back to control location when photoinhibition light was on (blue) or during control conditions with no photoinhibition (grey). Data from 9 new positions in 5 mice t-test, ** p<0.01.

### Late signals in M1 are necessary for motor task adaptation

Are these performance outcome signals in M1 important for behavior? Our data suggests a working hypothesis that layer 2-3 holds outcome signals which it then uses to influence motor commands, either directly or indirectly, in the layer 5 PT network to adapt future behavior. If this hypothesis is true, then disrupting the outcome signals in M1 should disrupt an animal’s ability to change its movements given a new task requirement. To test this, we used a task variant where animals are forced to learn a new pellet position. In control conditions, moving the pellet away from the set position initially adversely affects success rates, but then adjustments to the arm trajectory are made, and skillful performance is partially restored within the same session (Figure 7E, G). To perturb the memory of the last trial’s outcome, we optogenetically silenced M1 cortex during the inter-trial interval (ChR2 activation of resident inhibitory neurons in *Gad2-Cre* mice or stGtACR2 (Mahn et al., 2018) silencing of pyramidal neurons in *Slc17a7-IRES-Cre* mice, after execution of most movements or movement attempts ended, 800-1000 msec after the tone). We find that this post movement cortical disruptions prevented learning the new pellet position (Figure 7F-G; success rate in new position 27.5 ± 13.9 % and 5.8 ± 4.8 % in control and optogenetic silencing respectively, p<0.01; 9 new positions in 5 mice). This result was not due to a direct effect on basic motor execution of the forepaw for two reasons. First, silencing was delayed until the post-reach epoch, allowing the initial motor act to be performed in an unperturbed manner. Secondly, in control experiments when the pellet remained in the original location, silencing cortex late in the trial, did not significantly change success rates (success rate 63 ± 15.3 % and 61 ± 10.6 % with and without optogenetic activation, respectively; 2 animals in 5 sessions).

These findings are consistent with the role of motor cortex in conveying outcome information crucial for motor learning and adaptive behavior.

## Discussion

In this study, we examine the role of two main cortical layers in M1 in outcome evaluation of skilled dexterous behavior. While PT neurons are mostly concerned with movement related activity during the trial, a major component of layer 2-3 activity is devoted to monitoring positive and negative motor outcome information. In layer 2-3, we find two neuronal populations, success and failure related cells, whose activity distinguishes the end-result of the trial. Outcome related signaling is not innate to the M1 layer 2-3 network, but rather develops with learning of the task. The performance outcome activity of these neurons is not representing reward per se but rather reflects a measure of the hand reach performance itself. We further show that a memory trace of the outcome history is contained in the initial state of PT network activity on the next trial. Our optogenetic experiments suggest that M1 cortical activity between behavioral attempts, which include both the performance outcome signals in layer 2-3 and subsequent effects on the layer 5 PT network, is necessary for animals to adapt to changes in task requirements. The outcome signals we report here, may be considered as higher level signals reporting the end result which may have relevance in correcting future movements but have no relevance to immediate movements. This type of reinforcement outcome signals can serve for motor learning and adaptation assigning credit or punishment in the process of learning over longer time scales as discussed below (Even-Chen et al., 2017; Schultz, 2000; Wolpert et al., 2011) and for maintaining learned motor skills (Uehara et al., 2019).

In principle, this success and failure related neuronal activity might result from kinematic differences between success and failure trials. However, the lack of correlations between specific kinematic parameters and success and failure signals in layer 2-3 neurons shown by our GLM analysis does not support such possibility. In addition, it is important to stress that the outcome signals we report here are not reporting success and failure of every single grab attempt, rather they develop at the end of the trial after the mice made several failed attempts before a successful grab or towards the end of multiple grab attempts in failure trials. Thus, these outcome signals cannot be simply related to specific kinematics movements of the hand but rather likely convey the outcome of the whole sequence in a trial. Although we cannot completely rule out differences in general posture or other minor body differences after success and failure trials, such differences if they exist do not detract from the main findings. If the animals post-trial behavior reflects the outcome of the last trial, something in the nervous system must be commanding this response. Therefore, the signals we find in cortex could be the signals remembering the outcome and commanding the post-trial behavior, they could be an efference copy from some other brain region commanding this behavior, or they could be reacting to the sensory consequences of the post-trial behavior. In all of these cases, the signals in cortex would still contain information about the outcome of the last trial.

In contrast to previous recent studies in primate limb motor cortex that reported only negative error or lack of reward signals (Even-Chen et al., 2017; Inoue et al., 2016; Ramkumar et al., 2016), here we report both positive and negative performance outcome signals. The reason for the discrepancies between our findings and those previously reported, may be related to several potential factors. The smaller number of neurons recorded in primate motor cortex; a possible selection bias towards larger neurons with extracellular recordings; or differences in the tasks. While tasks in primates involved adaptation in reaching guided by changing visual goals, in our case the task involves complex reaching and grabbing movements that results in rich sensory differences between success and failure trials. Finally, the lack of positive outcome signals may be caused by differences between primates and rodent motor cortex. This last possibility is less likely as positive outcome signals were reported in supplementary eye field (SEF) neurons in primates (Sajad et al., 2019; Stuphorn et al., 2000). It is interesting to note that positive outcome signals were also reported in ALM and vibrissae motor cortex, during a localization whisking task in mice (Chen et al., 2017). The layer representation of the signals we describe in M1 differed from those reported in both these specialized motor regions (SEF and vibrissae motor cortex). In contrast to our findings, signals in ALM and vibrissae motor cortex were more prevalent in layer 5 compared to layer 2-3 (Chen et al., 2017). In SEF, signals were distributed across all layers with negative reporting signals (loss of reward and errors) more prevalent in layers 2-3 (Sajad et al., 2019). These differences may reflect differences in the degrees of freedom of the effectors being controlled by each region. The presumably more complex control of the forelimb may necessitate separation of monitoring and control functions.

How are performance outcome signals generated in M1? These signals could be generated through “predictive coding” (Friston, 2011; Kording and Wolpert, 2006) where copies of motor commands could be used to generate a prediction of the end result of that action. This prediction could then be compared with the actual sensory consequence once the trial is executed (Izawa and Shadmehr, 2011; Wolpert et al., 2011). If a match exists between the prediction and the actual, a success signal could be generated. In contrast, if a mismatch is detected, a failure signal would be generated. If this is the underlying mechanism for the signals seen here, it will be of great interest to see where the prediction is being produced and where the comparisons are being performed. One possible source contributing to prediction generation, might be the cerebellum which is a major input source to M1 via the thalamus (Bosch-Bouju et al., 2013) and has been reported to generate diverse signals including outcome signals, reward delivery, reward omission and expectations (Heffley et al., 2018; Kostadinov et al., 2019; Wagner et al., 2017). Interestingly, in the cerebellum positive outcome signals were reported and are reminiscent to the success related activity we report here (Heffley et al., 2018). However, in contrast to the success related signals we describe here, which lasts many seconds, the time course of the responses at the Purkinje neuron dendrites last several hundreds of milliseconds (∼500-600 ms) only. In addition, success signals in Purkinje cells were also seen when reward was delivered outside the context of the behavior, which was not seen in M1 in the “tongue reach” experiments. Thus even if the source of the success related signal is driven by the cerebellum, this signal undergoes significant processing at M1 network. Of course the reverse possibility also exists that the outcome signals in cerebellum are dependent on M1 inputs. Other input sources such as S1 or VTA may also contribute to outcome signals in M1 (Hosp et al., 2011; Lacefield et al., 2019; Mao et al., 2011; Mathis et al., 2017; Molina-Luna et al., 2009; Schultz, 2000; Wickens et al., 2003).

An alternative but related mechanism for generating these signals is that sensory information alone creates two different activity states for success and failure trials. Such sensory-driven divergences in activity could be due to network modifications that arise from plasticity mechanisms during the learning process. Success and failure consequences are essentially cached, or stored in memory. In this scenario, online predictions built off the outgoing motor command are not necessary for creating the success and failure related signals. Distinguishing between predictive coding and a cached system would require separately manipulating sensory reafference or the copies of motor commands associated with success and failure trials.

We show that the initial state of the PT dynamical system on the next trial is related to the outcome of the previous trial. Although the initial state of the PT neuronal activity may not be directly caused by the success and failure related activity of layer 2-3 neurons, it is conceivable that the outcome activity in layer 2-3 contributes to the initial state of PT neurons as layer 2-3 neurons feed its information in almost unidirectional manner to layer 5 neurons of M1 (Weiler et al. 2008). Success and failure signals in layer 2-3 might induce distinct effects on inter-trial network activity in layer 5, either directly or indirectly via other brain regions such as premotor cortex, cerebellum or basal ganglia. For example, success signals may feed onto the PT network to promote network states similar to the previous successful trial. In contrast, failure signals may promote network states that will generate behavioral variation on the next trial. If failure signals are generic error messages, the effects could be to put the PT network into a state that allows more explorations on future trials, analogous to a strategy thought to be used in songbird learning (Kojima et al., 2018). On the other hand, if the failure outcome signals provide information about the nature of the error, they might influence the network to produce a more specific adaptive correction. Further work is needed to firmly establish the link and distinguish between the different possible mechanisms that could link performance outcome signals and the change in the activity state of PT neurons and behavioral changes on subsequent trials.

Our optogenetic experiments show that M1 cortex activity late in the trial is needed for adapting future behaviors to changes in task requirements. These findings are consistent with a role of outcome signaling in motor adaptation in light of the temporal overlap between the activation dynamics of success and failure related neurons and our optogenetic mediated network inhibition. However, a clear causal relationship cannot be proven by these experiments as we did not silence selectively the success and failure related neurons. Silencing selectively the success and failure related neuronal population poses an experimental challenge, and further methodological developments are needed to achieve this goal. It should be stressed that memory of outcome information in the inter-trial interval could be stored in other brain regions, but our results suggest that this memory still depends on normal M1 activity. Interestingly, similar perturbations during well learned behavior do not have an effect on task performance on the time scale of our experimental paradigm (∼ up to 60-80 trials). This result indicates that the outcome signals we describe here, do not serve as an immediate trial by trial corrective error signals such as the signals described in Inoue et al. Rather, as these signals report end result of trials, they may serve as a reinforcement signals for learning over longer time scales. Our experiments were performed on trained animals that nearly plateaued their performance such that further improvement of task execution can only be achieved over long time scales. In this plateaued performance, ‘success’ signaling does not easily appear to drive further improvement of task execution and may be used to maintain the learned motor program. The absence of this maintenance signal may begin to degrade the model, but this might occur over longer time scales than those tested in our experimental paradigm. Moreover, here, cortical perturbations were done after both success and failure attempts, and therefore we have affected both outcome signals. The absence of either signal may lead to less circuit effects and consequently less effects on performance over short time scales.

Why should M1 use performance outcome signals over reward signals in the context of reinforcement learning? An important advantage of a performance outcome signal is that it is necessarily linked to the specific movement being learned. This is not the case for reward based signaling. Animals can acquire the same rewards using many different kinds of movements and environmental settings (Schultz, 2000). If an animal did use another effector (i.e. the tongue) to acquire the reward and a global reward signal was used, it could affect the stored program for the original effector. If the animal licked for the reward, the efference and reafference arm signals would be absent and other spurious inputs might be present. This could cause non-pertinent information to be incorporated, leading to degradation of the arm motor program. The absence of success and failure related signals during tongue reach and the presence of outcome signals in the plastic pellet tasks make a strong case that such interference is not an issue in M1. By being necessarily linked to the efference and reafference signals, performance based signals can ensure that modification to synapses or circuit dynamics are specific to circuitry relevant to the stored motor program.

Subpopulations of neurons serving distinct functions is a fundamental concept in neural networks (Adesnik and Naka, 2018; Zeng and Sanes, 2017). Layer 2-3 neurons dramatically differ from layer 5 PT neurons in their input-output pattern, connectivity and intrinsic electrophysiological properties (Anderson et al., 2010; Harris and Shepherd, 2015; Hooks et al., 2011; Tsubo et al., 2013), thus it is not surprising that these different neuronal populations may perform different computations in motor cortex. Previous work has shown activity differences between layer 2-3 and layer 5 neurons (Chen et al., 2017; Heindorf et al., 2018; Huber et al., 2012; Isomura et al., 2009; Komiyama et al., 2010; Masamizu et al., 2014), but here we report a new role of layer 2-3 neurons in generating performance outcome related signals during dexterous movements. An interesting hypothesis is that the observed separation of outcome evaluation (layer 2-3) and movement generation (layer 5) is beneficial in some way. Perhaps this is because the system needs to have an evaluation network not subjected to history effects, while such history effects are adaptive when employed in the movement command centers. Separation might also allow different plasticity rules to be operating in the different networks (Raymond and Medina, 2018). Interestingly, artificial deep neural networks use layer separation to increase computational efficacy.

In conclusion, positive and negative outcome signals exist within subnetworks of motor cortex, potentially allowing cortex to evaluate the past in order to adapt future behavior. The capacity to utilize performance signals may be a key reason why motor cortex is so essential for skilled dexterous behaviors (Whishaw, 2000). How outcome signals teach the pattern generators and forward models remain unanswered (Wolpert and Miall, 1996), but identifying a locus of representation of outcome signals is a necessary first step.

## Figure legends

**Figure S1.**
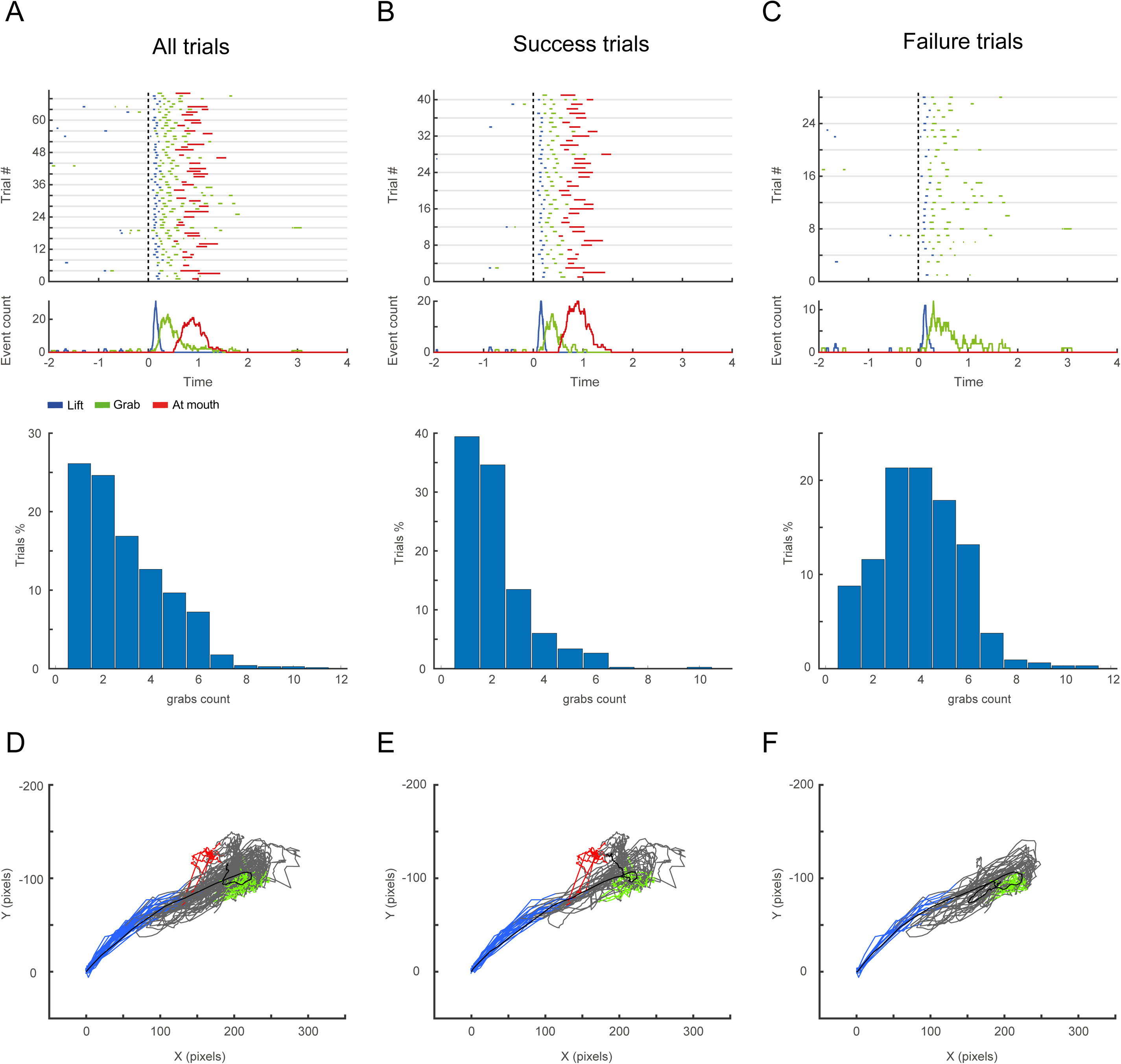
Behavioral ethograms. **A**. An ethogram annotating the behavior of an expert mouse over consecutive trials, performed by the modified JAABA software. Blue, hand lift; green, grab; red, food at mouth; dashed line represents tone. Lower panel, distribution of the number of grabs during the same session. **B**. Same as in A, for success trials. **C**. Same as in A, for failure trials. **D**. Forepaw trajectories (side view) performed with DeepLabCut. Behavioral events are overlaid on the trajectories (Blue, hand lift; green, grab; red, food at mouth). Same session as in A. **E**. Same as in D, for success trials. **F**. Same as in D for failure trials.

**Figure S2.**
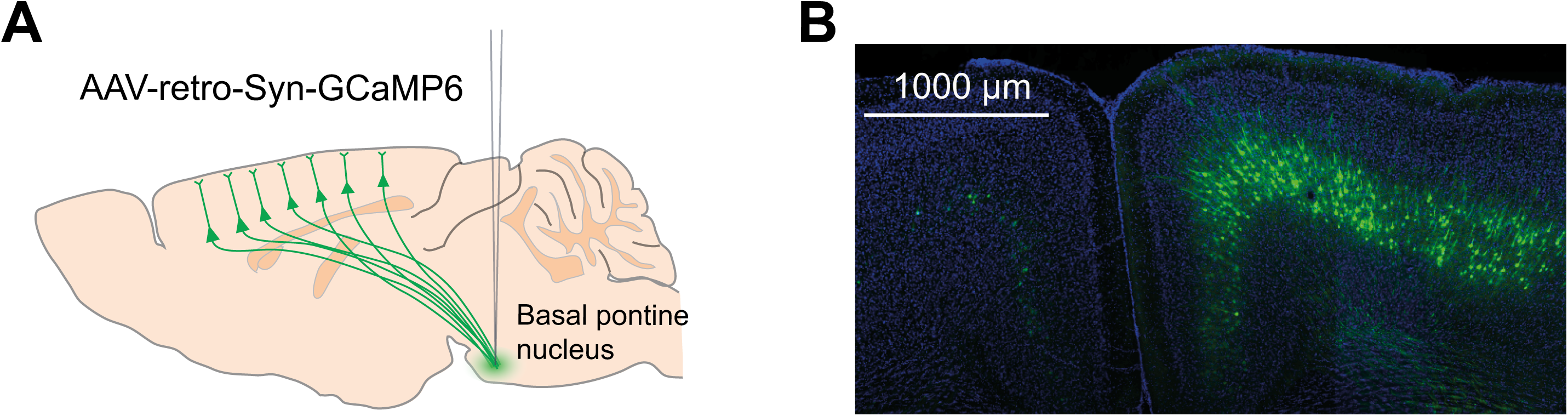
rAAV2-retro expression in layer 5 PT neurons in M1. **A**. Schematic of the retrograde injection paradigm. To label PT neurons, rAAV2-retro-GCaMP6 was injected to the basal pontine nucleus (coordinates 4.0 mm posterior and 0.4 mm lateral to Bregma, depth 5.4, 5.6 and 5.8 mm). **B**. Histology of an injected brain (coronal section) showing GCaMP6 expression in PT neurons (green) and DAPI staining to visualize cell bodies (blue).

**Figure S3.**
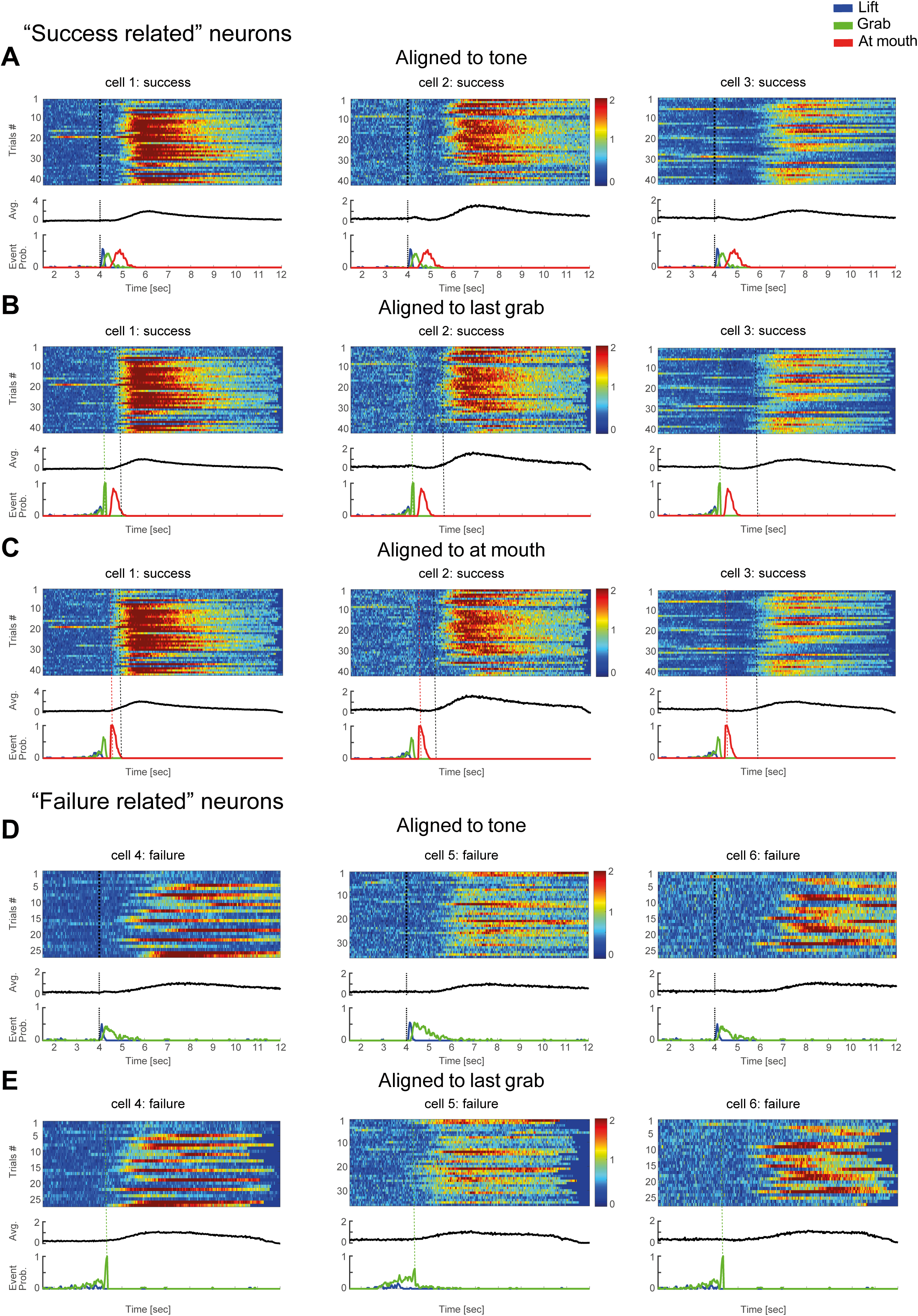
Example “success” and “failure” related neurons aligned to different behavioral and trial events. Examples of neurons (same neurons as in Figure 2A), aligned to tone, last grab and at mouth. All success (42 trials) or failure (28 trials) in one experimental session are presented for a single neuron. Each row shows ΔF/F of a single trial (color denotes the amplitude) aligned to one of the three events: **A**. Success related neurons aligned to tone. **B**. Success related neurons aligned to last grab. **C**. Success related neurons aligned to at mouth. **D**. Failure related neurons aligned to tone. **E**. Failure related neurons aligned to last grab. In all panels the black trace is the average over the presented trials; below, event-probability distribution for tone (black), lift (blue), grab (green) and at mouth (red). The dashed colored vertical lines demarcate the time of the aligned event (color according to the event). The onset of the calcium transients in success trials is denoted by grey dashed vertical line.

**Figure S4.**
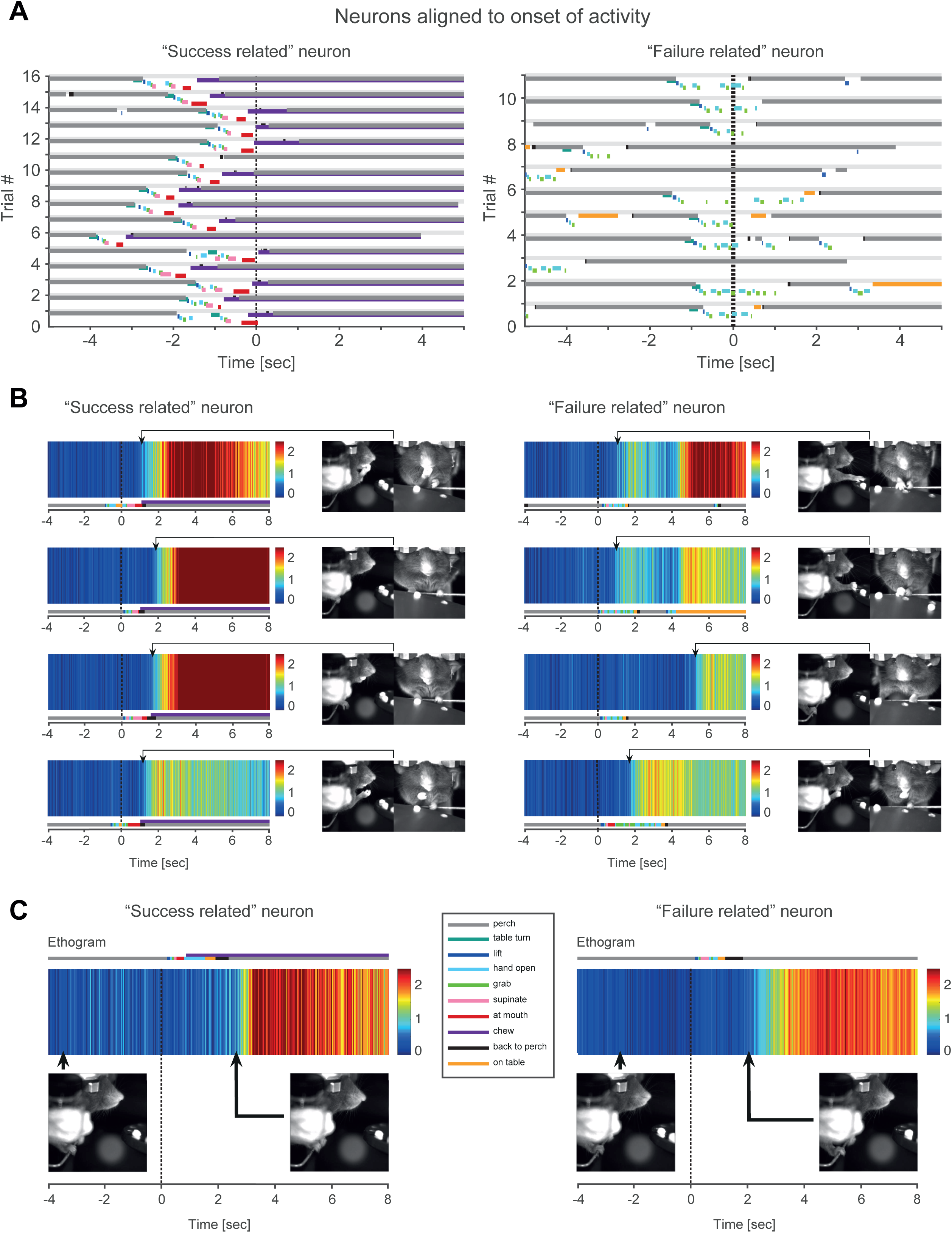
Alignment of behavioral events to calcium activity onset. Onset of activity of individual success and failure neurons could occur during different motor behaviors across attempts. **A**. Examples of alignment of the behavioral ethograms to the onset of success or failure related activity (denoted as “0”, dashed black line) of a single failure (right panel) and success (left panel) neuron. Onset of calcium transients was determined when the ΔF/F reached a value of 3SD above the mean. Behavioral events are color coded (see legend). **B**. Examples of single trials from a failure related (right panel) and a success related (left panel) neuron showing the calcium transient (upper panel), the behavioral events (lower trace). The time of activity onset is indicated by the black arrow head and the corresponding behavior at that time frame is shown on the right. **C**. Example of two trials from a failure (right panel) and success related (left panel) neurons taken from the same experimental session, showing failure and success signals during same behavior (hands on perch). Colored bar shows the color coded behavioral events. Note the similarity in the behavior before the cue onset and after hand movement ended (snap shots are shown below for each trial), the success or failure related signal appears only in the context of the behavioral outcome, either success or failure.

**Figure S5.**
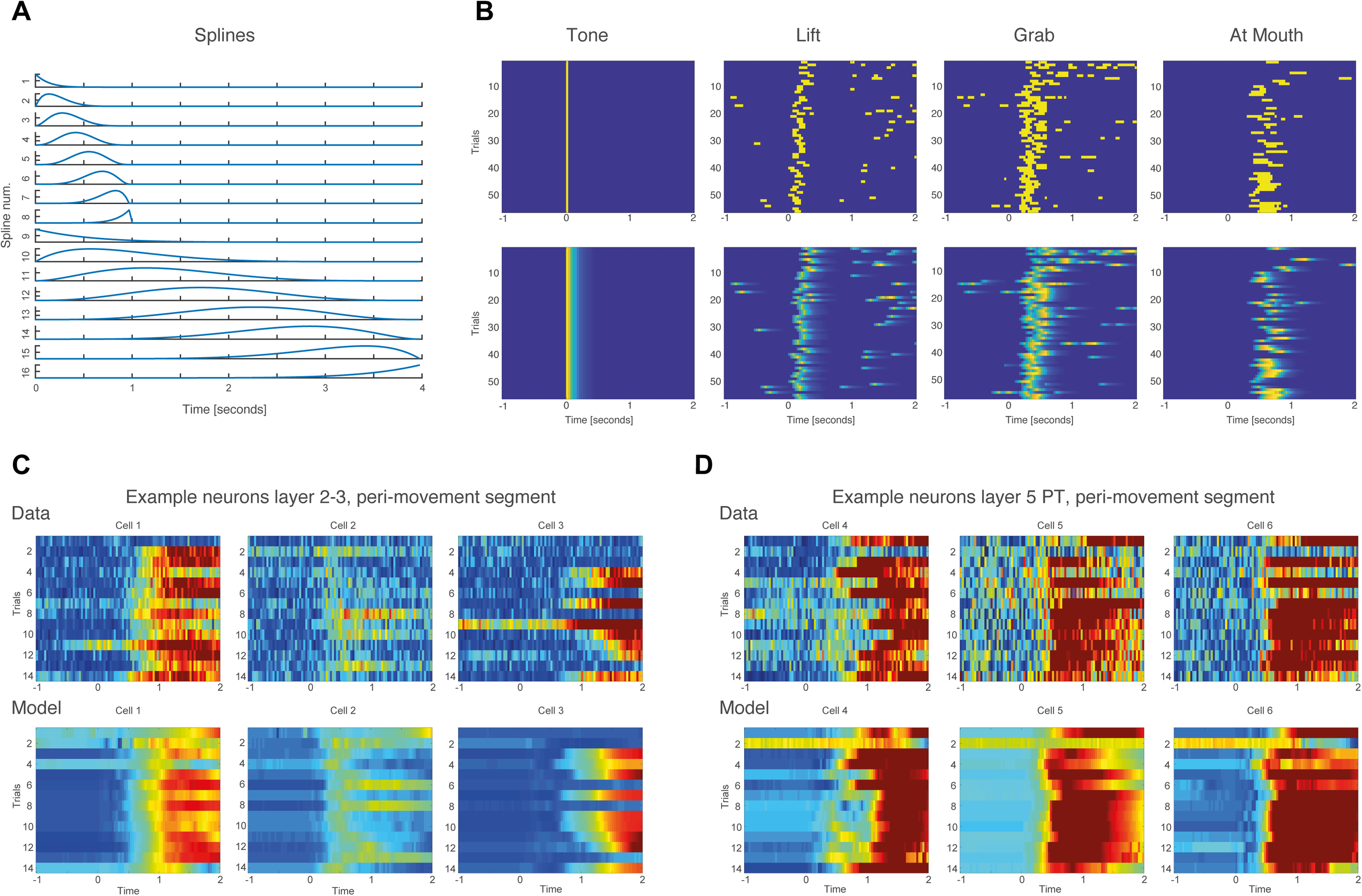
Modeling single neurons calcium transients with GLM. **A**. To model the time course of calcium transients a set of 16 splines with fast (0.5 sec) and slow (2 sec) time course were generated. **B**. The splines were convolved with the different predictors tone, lift, grab and at mouth. Examples are shown for one session, peri-movement segment (Tone -1 to Tone +2 sec). Upper panels are the annotated predictors and lower panels are the convolved predictors with the splines. **C**. Examples of modeled neurons in the peri-movement segment. Three neuron examples for layer 2-3. **D**. Three neuron examples for layer 5 PT neurons. Upper panels, experimental data; lower panels, modeled activity.

**Figure S6.**
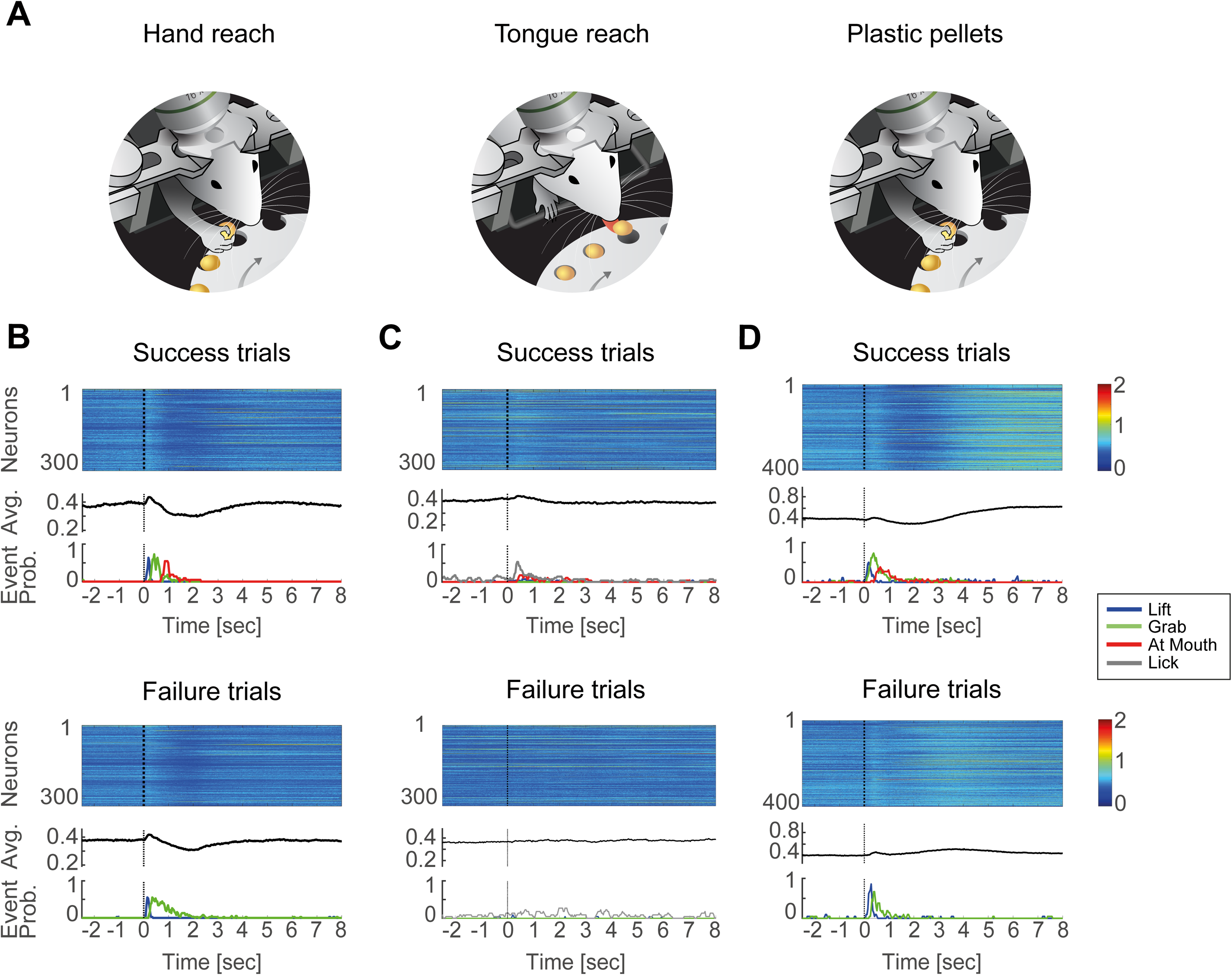
Average cortical activity for hand reach, tongue reach, and hand reach with plastic pellets. **A**. Mice were trained in different experimental conditions either hand reach with normal pellets (left panel), tongue reach with normal pellets (middle panel) or hand reach with plastic pellets (right panel). **B**. Upper panel, averaged calcium transients for all layer 2-3 pyramidal neurons over success trials in a single session during consecutive hand reach trials. Color encodes the percent change in fluorescence (ΔF/F). Black trace, grand average over all neurons and trials. Event-probability histograms for the behavioral events lift (blue), grab (green), at mouth (red), lick (grey) presented aligned in time. Dashed black line denotes the time of the tone. Lower panel, for failure trials. **C**. Same as in B for tongue reach. **D**. Same as in B for hand reach with plastic pellets.

**Figure S7.**
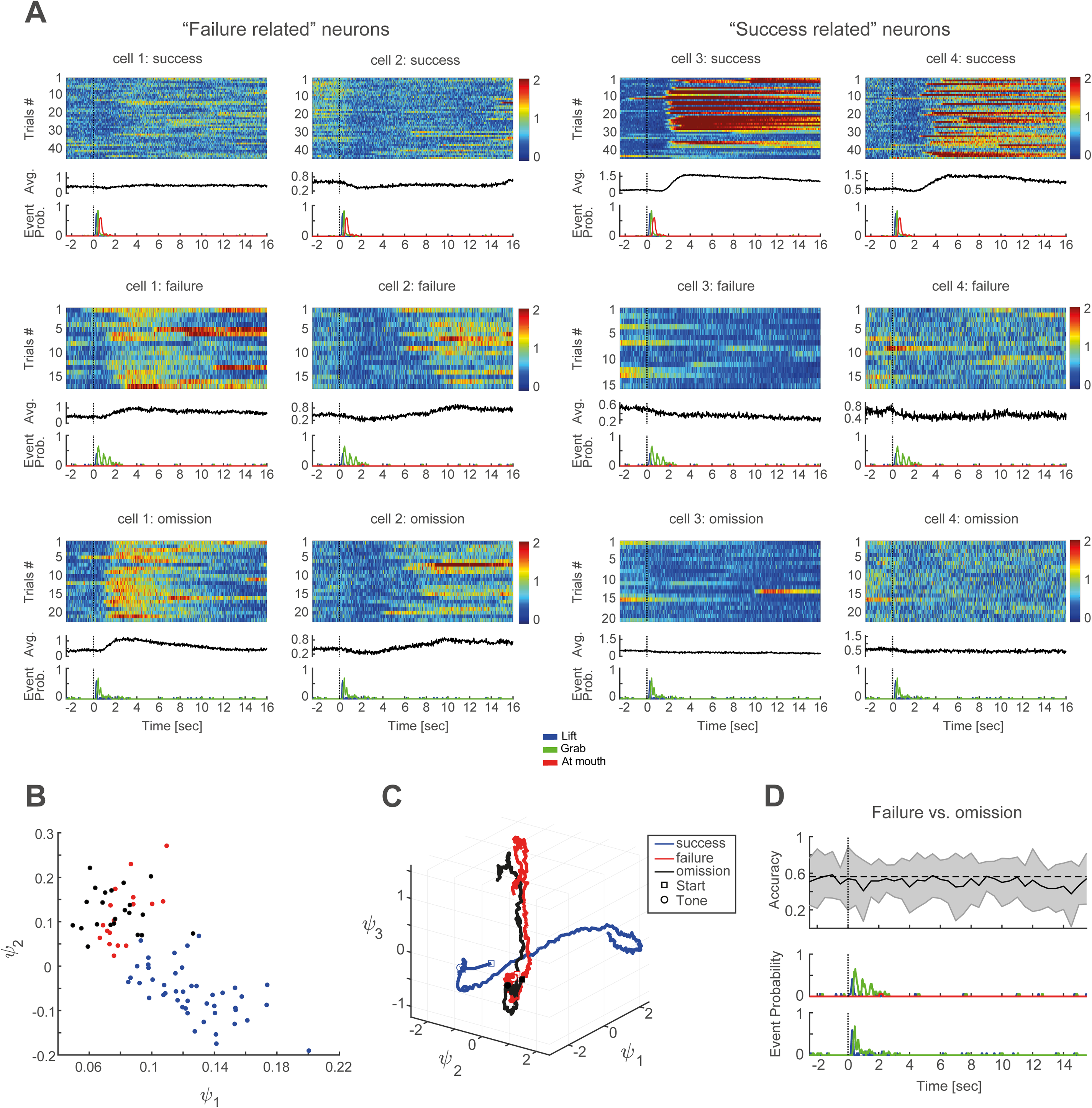
Cortical activity during omission trials is similar to failure trials. **A**. Examples of L2/3 failure related neurons (cells 1 and 2) and success related neurons (cells 3 and 4). Activity is presented over success (upper panel), failure (middle panel) and omission (lower panel, omission 26% of trials) trials during one experimental session. **B**. 2D PCA embedding for layer 2-3 neurons in one example experiment with omission trials (red, failure; blue, success; black, omission; same experiment as in A). Accuracy of separation between failure and omission trials was 0.56±0.23 and between success and failure trials was 0.97±0.07. **C**. Average temporal evolution of the network activity of layer 2-3 neurons separated to success (blue), failure (red) and omission (black) trials (same experiment as in A and B). Sensitivity index for success vs. failure was 20 and for failure vs. omission 9. **D**. SVM accuracy of separability for layer 2/3 activity in failure vs. omission trials (± SD shown in grey, dashed line represents chance level). Average accuracy 0.56, chance 0.5. Same experiment as in A, B and C.

**Movie S1.** Example of successful hand-reach with normal pellets. Side and front views captured with 200Hz cameras. After an auditory cue, pellet was delivered on a rotating table. The mouse grabbed the food pellet with its forepaw and consumed it. Playback at 60 Hz.

**Movie S2.** Example of successful tongue-each. Side and front views captured with 200Hz cameras. After an auditory cue, pellet was delivered on a rotating table. The mouse licked the food pellet with the tongue and consumed it. Playback at 60 Hz.

**Movie S3.** Example of successful hand-reach with 3D printed plastic pellets. Side and front views captured with 200Hz cameras. After an auditory cue, pellet was delivered on a rotating table. The mouse grabbed the food pellet, brought it to its mouth, and then discarded the pellet. Playback at 60 Hz.

**Movie S4.** Animation of averaged temporal trajectories of layer PT neuronal activity sorted according to outcome history. Average trajectories (3 first PCs) were sorted according to previous and present trial outcomes resulting in 4 types of trajectories: previous and present successes (S-S, blue); previous failure and present success, (F-S, cyan); previous success and present failure, (S-F, violet); previous and present failures (F-F, red). Square marker denotes start of trial; circle denotes the time of the tone (4 sec after start).

## Acknowledgments

We thank Britton Sauerbrei, Kimberly Ritola, James Phillips, Jihong Zheng, Jian-Zhong Guo, Kristin Branson and Blake Richards for helpful discussions throughout the project and helpful comments on the manuscript. We thank Shay Achvat for software assistance, Irena Reiter for technical assistance, Julia Kuhl for illustrations in the manuscript and Zarixia Zavala-Ruiz for helping with the Janelia visitors program. This study was partially supported by Janelia visitors program (JS), the BSF-NSF/NIH Foundation (JS, RT and RM), the MORASHA ISF Biomedical Research Program (JS and YS), Adelis Fund for Brain Research at the Technion and Prince funds (JS and YS).

## STAR★Methods

### KEY RESOURCES TABLE

| REAGENT or RESOURCE | SOURCE | IDENTIFIER |
| --- | --- | --- |
| Bacterial and Virus Strains |  |  |
| pAAV.Syn.Flex.GCaMP6s.WPRE.SV40 | Addgene | Addgene, 100845-AAV1 |
| AAV-retro-Syn-GCaMP6s | Tervo et al. 2016 | Janelia virus core |
| AAV1.CAGGS.Flex.ChR2-tdTomato.WPRE.SV40 | Penn vector Core | N/A |
| pAAV_hSyn1-SIO-stGtACR2-FusionRed | Mahn et al. 2018 | Addgene, 105677-AAV1 |
| Experimental Models: Organisms/Strains |  |  |
| Mouse: C57BL6 | ENVIGO | N/A |
| Mouse: Slc17a7-IRES-Cre | Janelia research campus HHMI | N/A |
| Mouse: GAD2-cre | Yizhar lab, Weizmann Institute of Science | N/A |
| Software and Algorithms |  |  |
| JAABA | Kabra et al. 2013 | http://jaaba.sourceforge.net/ |
| Matlab | MathWorks | http://mathworks.com |
| DeepLabCut | Mathis et al. 2018 | https://github.com/AlexEMG/DeepLabCut/ |
| LIBSVM | Chang and Lin 2011 | http://www.csie.ntu.edu.tw/~cjlin/libsvm |

### LEAD CONTACT AND MATERIALS AVAILABILITY

Further information and requests for resources and reagents should be directed to and will be fulfilled by the Lead Contact, Jackie Schiller.

### EXPERIMENTAL MODEL AND SUBJECT DETAILS

All animal procedures were in accordance with guidelines established by the NIH on the care and use of animals in research and were confirmed by the Technion and Janelia Research Campus Institutional Animal Care and Use Committee. Adult male C57BL6, Slc17a7-IRES-Cre and GAD2-cre mice were used in this study. Animals were housed in a 12:12 reverse light:dark cycle. For behavioral training and experiments food intake was limited to 2.5–3 g/day with ad libitum water.

### METHOD DETAILS

#### Experimental Design

To investigate outcome representation in M1 we performed two photon calcium imaging experiments in awake head-restrained mice performing a skilled forelimb reach and grasp task. Chronic calcium imaging was performed with the genetically encoded Ca2+ indicator, GCaMP6, that was introduced to the cells via a viral vector injection which caused permanent expression of the indicator, allowing us to monitor the activity for many weeks. We selectively labelled layer 2/3 pyramidal neurons or L5 PT neurons using Cre-dependent expression or a newly developed retrograde viral vector (Tervo et al. 2016), respectively. We implanted a chronic window over the M1 forelimb area that allowed us to image the activity for long time periods.

#### Viral injections and cranial window surgery

Male mice were housed in a 12:12 reverse light:dark cycle. Surgical procedures were performed under isoflurane anesthesia (4% for induction and 1.5-2% during surgery) at the age of 2-3 months. We performed a circular craniotomy (2.5 – 3 mm diameter) centered at primary motor cortex forepaw area (0.6 mm anterior and 1.6 mm lateral to Bregma (Silasi et al. 2013)). The imaging window was constructed from two layers of 170 µm thick panes of laser-cut glass glued together with an optical UV adhesive (Norland). A custom-made headpost (Osborne and Dudman, 2014) was affixed to the skull using dental cement. Layer 2/3 pyramidal cells were labelled by locally injecting AAV-Syn-flex-GCaMP6s to the left M1 forepaw area (∼300 µm depth, 60 nl) to Slc17a7-IRES-Cre mice during the window implantation surgery. To label L5 PT neurons, AAV-retro-Syn-GCaMP6s (Tervo et al. 2016) was injected to the left basal pontine nucleus (4.0 mm posterior and 0.4 mm lateral to Bregma, depth 5.4, 5.6 and 5.8 mm, 100 nl per site) of C57BL6 mice. The injections were made through the thinned skull using a hydraulic micromanipulator (M0-10 Narishige). Ketoprofen (5 mg/kg) and buprenorphine (0.1 mg/kg) were administered subcutaneously for analgesia during surgery and for 2 days post operational to reduce inflammation and analgesia respectively. Mice recovered for at least 1 week following surgery with ad libitum food and water.

#### Behavioral training

After recovering from surgery, animals were food restricted by limiting food intake to 2.5–3 g/day with ad libitum water. Training started when the animals reached 85-90% of their original body weight. Mice were habituated to head fixation in a custom-built apparatus (Osborne and Dudman, 2014) in dark and quiet conditions, monitored by a webcam. Mice were initially trained to retrieve food pellets (14-20 mg; Test Diet; St Louis, MO) from a rotating plate placed directly below their mouth. The plate was rotated every 60 sec using a servomotor (Gecko Drive) driven by custom-made Arduino software (200 msec duration) to present a food pellet. An auditory tone (200 msec, 1 kHz) was used as a cue during plate rotation. Mice were trained daily for 20-30 min until they routinely responded to the auditory cue and grabbed the food pellet with at least 50% success, thereafter behavior was combined with two-photon imaging. Care was taken to reproduce the pellet location every session for every animal. Animals typically succeeded retrieving the food pellet using their forepaw for the first time after 3-5 training sessions. Mastering the task typically took approximately another 10 sessions.

For tongue reach experiments, expert hand reach task mice were retrained to access the food pellet with their tongue.

#### Two-photon calcium imaging

Images (512 × 512 pixels) were acquired at 30 Hz using a two-photon microscope equipped with a resonant scanner (resonant scanner frequency 8kHz; Brucker Corp. and custom made Janelia scope), 16X water immersion lens (Nikon, 0.8 NA, 3 mm working distance) and controlled by the software package PrairieView 5.3 or Scanimage. GCaMP6s was excited at 940 nm using a femtosecond pulsed laser (InSight X3, Spectraphysics or Coherent Chameleon Ultra II). Emission light was detected by a GaAsP photomultiplier tube (Hamamatsu).

Each trial consisted of 12 sec total duration, with the tone and plate rotation introduced at 4 sec from trial start (Figure 1). Inter-trial intervals were typically 30 sec and a total of 50 - 120 trials were collected per single experimental session. The same field of view was imaged over all experimental sessions of the same animal. Behavioral performance was monitored at 200 Hz using two cameras (side and front view; Flea3 FL3-U3-13Y3M, PointGrey). The time coordination of two photon calcium imaging, behavioral task and video recording was accomplished via a National Instruments board (PCI-6110) using custom made software written in Matlab.

#### Optogenetic silencing of M1

We used two methods to silence post movement activity of M1: 1. We injected AAV1CAGGSFlexChR2-tdTomatoWPRESV40 (Penn vector Core) to M1 (0.6 mm anterior and 1.6 mm lateral to Bregma, 60 nl) in GAD2-cre mice (n=3 mice) to activate resident interneurons. 2. We injected pAAVhSyn1SIOstGtACR2FusionRed (Addgene; Mahn et al. 2018) to M1 in Slc17a7-IRES-Cre mice (n=2 mice) to silence preferentially pyramidal cell somata. A chronic window was placed over the injection area, a head post was affixed to the skull and animals were trained in the forepaw reach and grab task as previously described. To selectively silence the late activity without affecting the forepaw movement we first performed regular reach and grab task experiments and calculated the average time when most hand movements ended for each animal (*T*_*e*_, usually ∼800 – 1000 msec after the tone). Optogenetic stimulation started at the calculated *T*_*e*_ for each animal and continued until the end of the trial. For ChR2 activation, a train of 10msec pulses was given at 10 Hz, using a 470 nm LED driven by a LEDD1B LED driver (ThorLabs) focused through a 16X water immersion lens (Nikon, 0.8 NA, 3 mm working distance), light intensity was 10 mW as measured coming out of the objective. For stGtACR2 silencing, one continuous pulse was given using a 447nm laser (OEM Laser Systems, Utah, USA) connected to an optic fiber placed over the window area, laser intensity was 4-8 mW as measured at the tip of the fiber. A blue LED was continuously lit in the vicinity of the animal to mask any light from laser activation. Control of the laser, behavioral task and cameras was achieved using custom routines written in Matlab. Both methods showed similar results and data were pooled for analysis.

#### Histology

At the end of experiments, the animals were deeply anaesthetized and transcardially perfused with 0.1 PBS followed by a solution of 4% paraformaldehyde. The brains were removed and stored in the fixative for 24-48 hours. Coronal sections were made at 50-100 µm thickness. The sections were mounted on slides embedded in Vectashield Hard Set mounting medium containing DAPI (Vector Laboratories) and imaged using a Pannoramic slide scanner (3D Histech).

### QUANTIFICATION AND STATISTICAL ANALYSIS

#### Behavioral data analysis

We used a modified version of the Janelia Automatic Animal Behavior Annotator (JAABA) software package (Kabra et al. 2013) to classify behavioral events. The hand reach task was segmented into discrete behavioral events (Lift, Grab and AtMouth). A subset of trials was manually labelled to train the classifier to recognize the behavioral events of interest. Then, machine learning based automatic classification was performed and behavioral events were extracted from all trials. The events were defined as follows: Lift was defined from initial separation between hand and perch until the hand reached maximum height, Grab was defined from the beginning of finger closure until the hand was lifted off the table (with or without the pellet), AtMouth was defined as hand with pellet in close proximity to mouth.

In the tongue reach task, we added another behavioral tag, Lick, which was defined when the tongue was first visible out of the mouth and until it returned inside the mouth, AtMouth was defined from when the food pellet reached the lips until the animal started to chew.

Success trials were defined as trials where mice succeeded in grabbing and bringing the food pellet to the mouth for consumption (regardless of how many grab attempts were made). Failure trials were defined as trials where mice attempted to grab but missed the food pellet and thus did not consume the food pellet. For tongue reach experiments, success trials were defined as trials where animals licked for the food pellet and managed to bring it to the mouth for consumption (regardless of how many lick attempts were made), and failure trials were defined as trials where animals licked for the pellet but did not manage to successfully retrieve it to the mouth and consume it.

We also used a machine-learning-based algorithm (DeepLabCut, Mathis et al. 2018) to automatically track the hand position in our behavioral videos. We manually labeled the position of the hand in a small subset of video frames to train the algorithm and then x and y locations of the hand were automatically extracted from each frame of the side and front videos.

#### Two photon data analysis

The fluorescence data acquired by the two photon microscope was first registered to correct for brain motion artifacts. Our registration method was based on (Kowalczyk 1990), using Fourier transform based correlation between two successive images. The maximal value position in the correlation image specifies the relative shift between the two images, we designate them *u*_*t*_ and *v*_*t*_. In this method a template specification and matching against an image stack is required. The template image *I*_*temp*_(*x, y*) was defined as the average of all images in the selected trial over time.

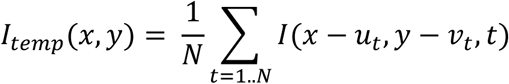

The set {*u*_*t*_, *v*_*t*_}, *t* = 1.. *N* is an image shifted in the XY plane after alignment. We initially start with *u*_*t*_*=0, v*_*t*_*=0* and then update their values according to the registration maxima. This procedure is repeated several times, when each time we compute the new template *I*_*temp*_(*x, y*) using previously computed {*u*_*t*_, *v*_*t*_} for each image. Typically, this procedure converges after several iterations, in our case 3 iterations.

To align the imaging data over many trials we used a similar technique, utilizing the previously computed averaged templates for each trial. For each trial *k*, we performed a single trial registration using the template algorithm for 3 iterations. To align the image data over many trials, we treated the final templates 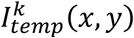 from each trial *k* as unaligned image data and repeated the same registration procedure to find offsets {*u*_*k*_, *v*_*k*_} for each trial. These offsets along with previously found offsets {*u*_*t*_, *v*_*t*_} account for the final image shift in XY plane.

Regions of interest (ROIs) were detected manually using average fluorescence images and ΔF/F projection images which highlighted active neurons. The pixels within each ROI were averaged for every frame. The ROI “mask” was used to detect the same neurons on multiple imaging sessions on different days.

*ΔF/F* was computed using the following formula:.

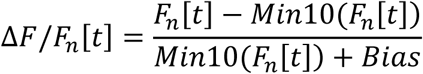

Where *Min10(Fn [t])* is a mean value of the lowest consecutive 10% values of the fluorescence signal *Fn [t]*. A small Bias factor in the denominator prevented zeros when the cell was completely silent.

#### Statistical Analysis

In each experiment, the mouse repeated the task for several dozens of times, performing a hand reach or a tongue reach task. The data were analyzed using a custom Matlab code, except for training and testing the SVM classifiers, for which we used the standard LIBSVM (Chang and Lin 2011).

The imaging data collected in each experiment are stored in a 3-dimensional matrix (tensor) **X** of size *N*_*r*_ ×*t* ×*T*, where *N*_*r*_ is the number of neurons, *t* is the number of time samples per trial, and *T* is the number of trials. Namely, *X* (*i, j, k*) is the neuronal activity of the i-th neuron at the j-th time sample in the k-th trial.

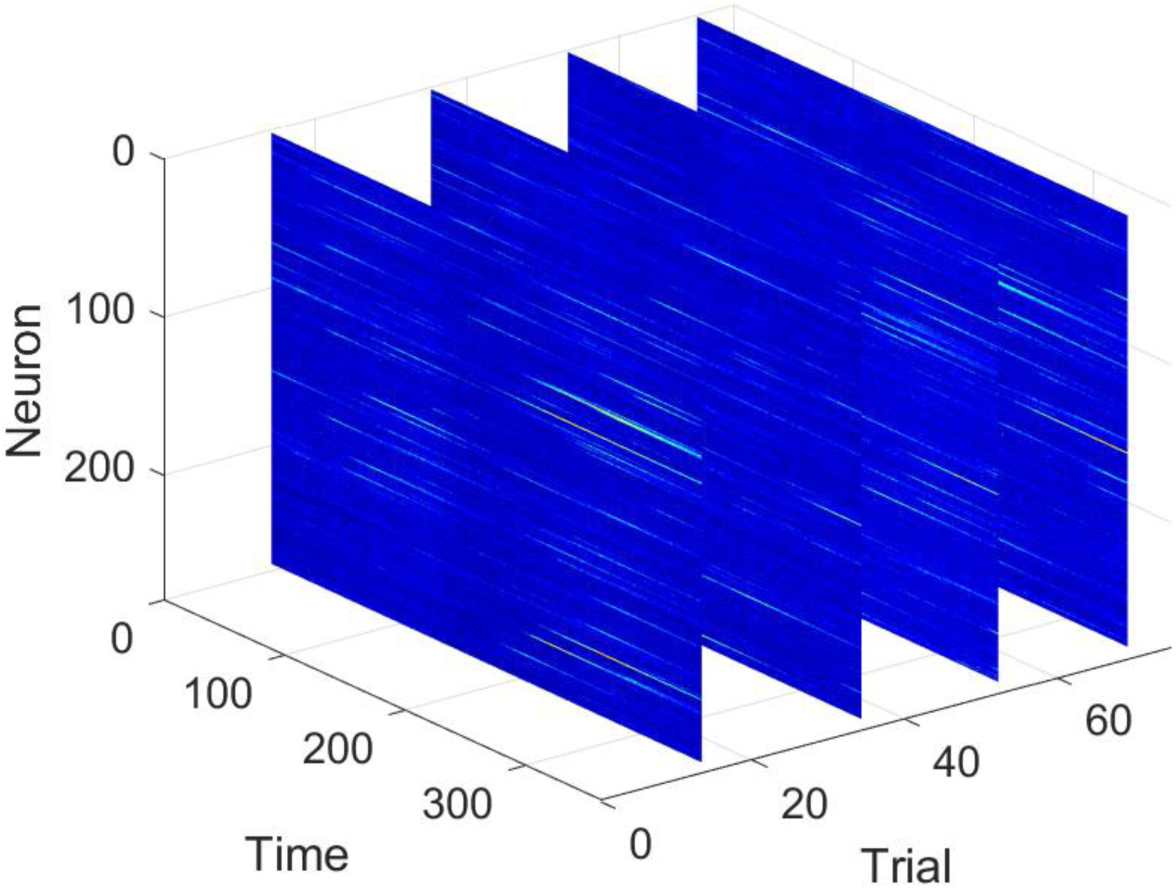

**An example showing the 3D matrix consisting of imaging data. Each slice consists of the neuronal activity over time of all neurons (ROI’s) in a particular trial**.

##### 2D PCA embedding of trials

The 3D imaging data matrix X includes the activity of hundreds of neurons over hundreds of time samples. Particularly, in order to distinguish between the activity of the network during success trials and the activity of the network during failure trials, we used Principal Component Analysis (PCA) to represent each trial in a lower dimensional space.

We defined the vector **x**_*i,k*_ as the temporal activity of the i-th neuron at the k-th trial, across time:

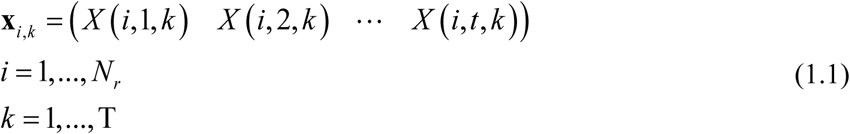

We reshaped the 3D imaging data matrix **X** as a 2D matrix of size *T* ×[*N*_*r*_*t*], where each row consists of all time samples from all the neurons related to a specific trial:

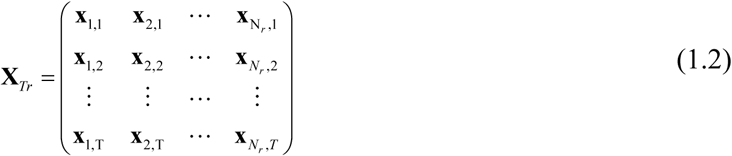

We evaluated the sample covariance of **X**_*Tr*_ by

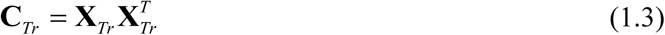

and applied the eigenvalue decomposition

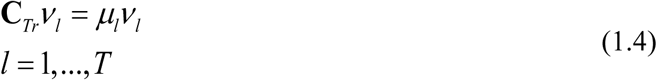

where *v*_*l*_ are the eigenvectors with the associated *μ*_*l*_ eigenvalues. The eigenvectors of the sample covariance give rise to the principal components of the imaging data tensor with respect to the trials axis.

We used the first two principal components of each trial, (*v* _*l*_ (1),*v* _*l*_ (2)), *l* = 1,…,*T*, to visualize the trials in a 2D Principal Component (PC) space, where the colors indicate the behavioral outcome (success – blue, failure - red) as depicted in Figure 4A and 4B.

##### Tree partition of trials using diffusion maps

In the previous section we demonstrated that a compact representation of trials, obtained by PCA, leads to a good separation with respect to the behavioral outcome of the trials. In this section we further examine the differences and similarities of trials, when nonlinearly embedded into a new Euclidean space and partitioned to a hierarchal tree using a new manifold learning method (Mishne et al. 2016).

In recent years, manifold learning has become a leading approach for detecting underlying structures and hidden parameters in data. Methods such as Laplacian eigenmaps (Belkin and Niyogi 2003), Hessian eigenmaps (Donoho and Grimes 2003) and diffusion maps (Coifman et al. 2005, Coifman and Lafon 2006) view the data samples as nodes of a weighted graph. The weights of the graph edges are determined based on some affinity measure, capturing the pairwise similarities and dissimilarities of the nodes, i.e. the data samples. Common practice is to represent the weights of the edges by a kernel. The eigenvalue decomposition of the kernel establishes a new Euclidean space, often of lower dimensionality, in which the data is embedded. Ideally, the embedding of the data in the new space reveals the main structures underlying the data.

Diffusion maps embed the data samples into a Euclidian space where pairwise distances describe the relationship between data samples in terms of their graph connectivity. This means that local structures of the data samples can be captured through their distances in the embedded space, which is the main reason why diffusion maps is broadly and successfully applied (Mishne et al. 2017, Shemesh et al. 2017, Yair and Talmon 2017, Shnitzer et al. 2016, Sulam et al. 2017).

In this work we used Diffusion maps where Euclidian distances between embedded data samples describe the relationship between them in terms of their graph connectivity (Coifman and Lafon 2006), which is just one of the reasons this method is used for many other data sources and computational purposes. Based on the matrix **X**_*Tr*_, which is of size *T* ×[*N*_*r*_*t*] and consists of the imaging data of each trial in its rows, we constructed a *T* ×*T* affinity matrix where each element is computed according to a non-linear function of the pairwise Euclidean distances between the trials,

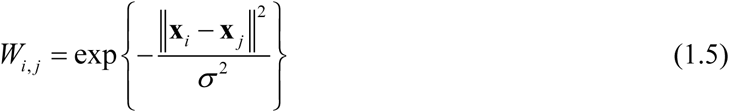

where *σ* is a scaling tunable parameter and **x**_*i*_ is the i-th row of 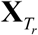. By normalizing the rows of **W**, we obtained a row stochastic matrix **P**,

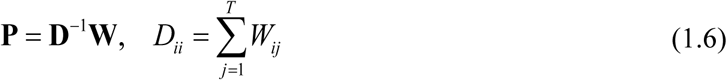

and computed its eigenvectors,

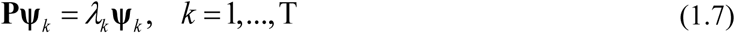

where **ψ**_*k*_ and *λ*_*k*_ are the eigenvectors and eigenvalues of **P**.

We defined Ψ_*k*_ as the diffusion map of the trials to a Euclidean space ℝ^*d*^ by

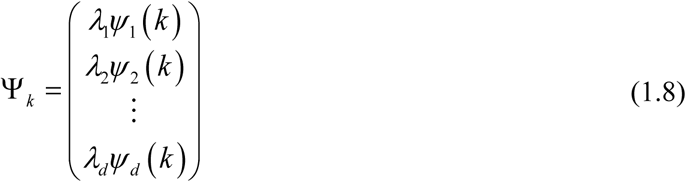

where *d* ≤ *T*. As shown in (Belkin and Niyogi, 2008), the matrix **P** can be viewed as a transition probabilities matrix of a Markov chain defined on the graph, whose nodes are the trials. The Euclidean distance between trials in the embedded space (1.7) constitutes the diffusion distance, which is related to aggregation of the transition probabilities on the respective graph.

We built a binary partition tree of the trials based on the diffusion distance, i.e. the Euclidean distance between their low dimensional representations in the embedded space (1.7). The construction of the partition tree is carried out in iterations in a top-down manner. In the first iteration, the embedded trials were partitioned into two clusters, which constitute the top level of the tree, just below the root. In the subsequent iterations, each cluster was further partitioned into two sub-clusters, forming the next level of the tree. These iterations were carried out until all the sub-clusters contained a minimal number of 12 trials. This particular number was set since it empirically led to good results in our experiments.

##### PCA trajectories

In this section we analyzed the dynamics of the network within each trial to demonstrate the differences between the temporal evolution of success and failure trials as well as the differences between Layer 2-3 and PT neurons.

We represent the temporal evolution of the entire network in each trial as a 3D trajectory using PCA, and therefore enable a more intuitive representation of the network’s dynamics. Focusing on the neurons axis, we reshaped the 3D imaging data matrix **X** as a 2D matrix of size *N*_*r*_ ×[*t* T], where each row consists of all the time samples from all the trials of a specific neuron as illustrated below.

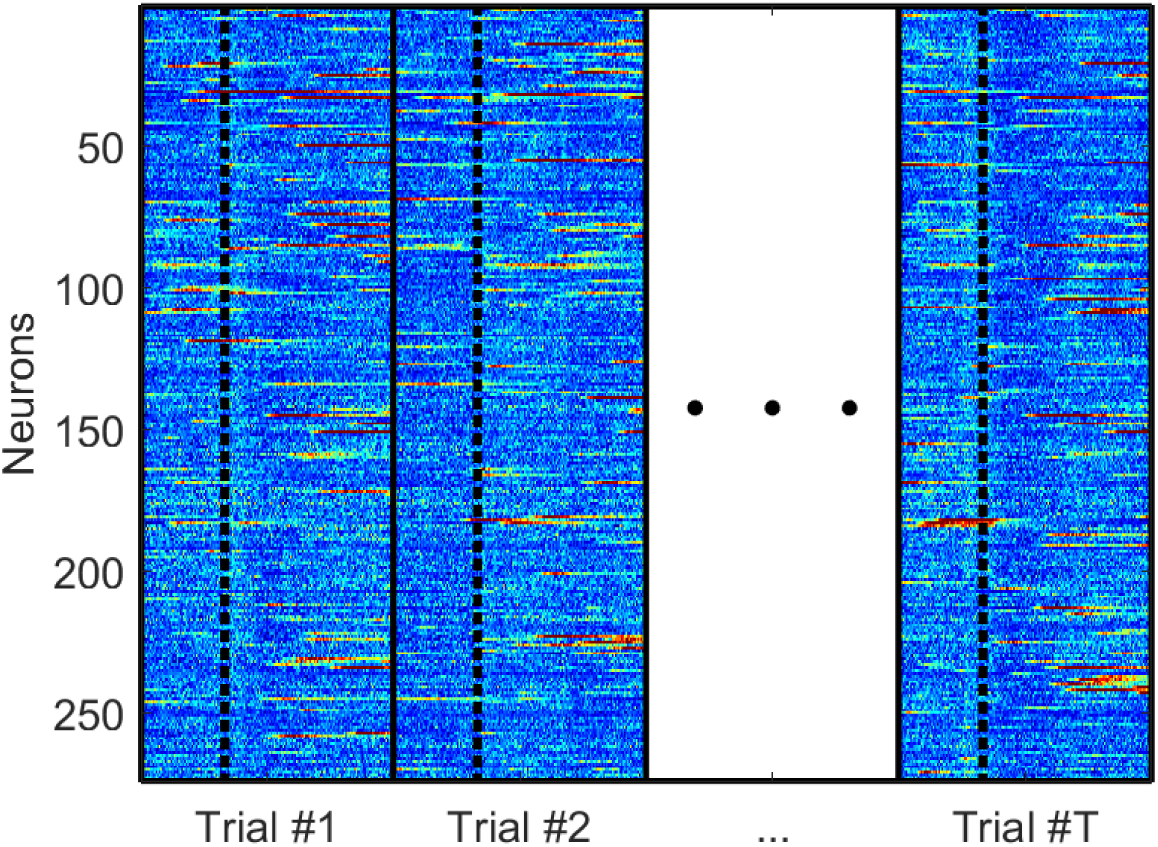

**Re-organization of the 3D imaging data matrix into a 2D matrix of size** *N*_*r*_ ×[*t* T]. **Borders of each trial are presented as solid lines and dashed lines demark the tone**.

Formally, similarly to (1.2), we denote this matrix by

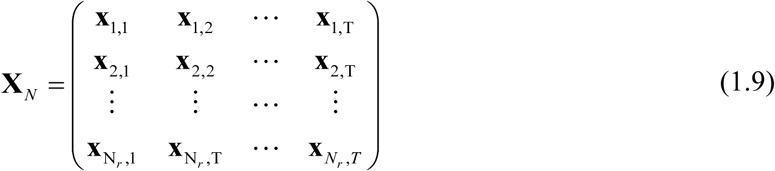

where **x**_*i*,k_ is defined in (1.1).

We computed the sample covariance of **X**_*N*_, denoted by **C**_*N*_, and applied eigenvalue decomposition to obtain its principal components **u**_*l*_,

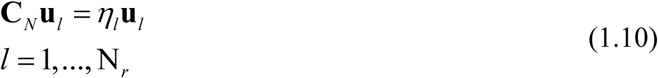

Projecting the imaging data onto the three principal components

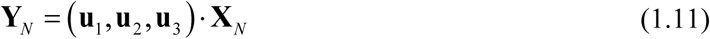

resulted in a 3×[*t* T] matrix, denoted by **Y**_*N*_. This matrix consists of a compact representation of the dynamic evolution of the network of neurons in a 3D coordinate system prescribed by the principal components. Explicitly, the elements of **Y**_*N*_ can be written as

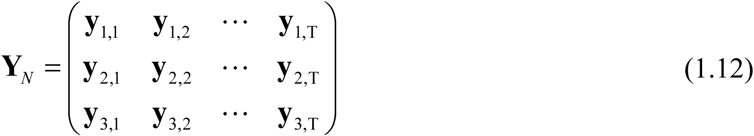

where **y**_*i,k*_ consists of the temporal activity of the network of neurons at the k-th trial, projected onto the i-th principal component.

Figure 4C and 4D present the temporal evolution of the network as 3-dimensional trajectories (fine lines) and the mean trajectories, averaged over the success and failure trials (thick lines, blue and red respectively).

##### Sensitivity index

To quantify the difference in terms of the dynamics of the network in the PCA space (1.10) between success trials and failure trials, we evaluated the sensitivity index (also known as d’ – d-prime) for each PC, i.e. **Y**_*N*_, as a function of time:

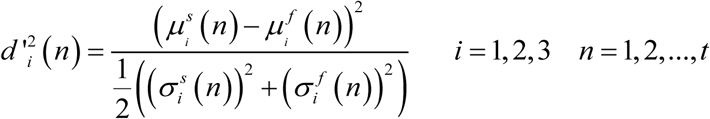

where 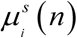 and 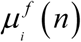 are the mean values of the i-th PC at time sample *n*, averaged over all success and failure trials, respectively. In the same manner, 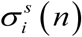 and 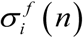 are the standard deviation (STD) values of the i-th PC at time sample *n*, associated with success and failure trials, respectively. The overall difference in terms of the dynamics between success and failure trials was obtained by the average value of the first 3 PCs, namely, by

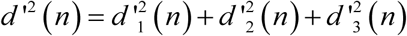

The sensitivity index as a function of time evaluated for layer 2-3 and PT neurons is presented in Figure 4E. It clearly indicates that the PCA trajectories indeed evolve differently through time, however, this separation is statistically more significant in Layer 2-3, than in PT neurons. The sensitivity index provides an objective evaluation for the success/failure separability, assuming an underlying statistical model consisting of two Gaussians representing the two classes. In the next section we describe another separability measure - the classification accuracy – which does not rely on this particular statistical model.

##### Classification Accuracy

To further quantify the difference between success and failure trials as conveyed by the dynamics of the neuronal activity, we use a linear SVM classifier (Joachims 2006). The classification accuracy was evaluated for each day of experiment separately. We used a sliding window of 1 second with 0.5 second hop. In every time window, we evaluated the average activity of each neuron at each trial,

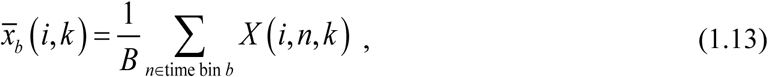

Where *b* is the index of time window and *B* is the window length measured in samples. The activity of the network in each time window for each trial was represented by the following *N*_*r*_ × 1 vector

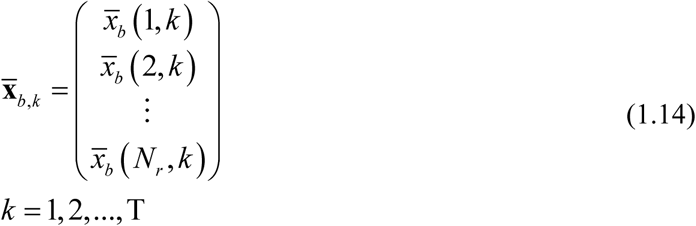

For each time bin, we paired the set 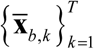, representing the averaged activity of the network across trials, with the associated success and failure labels,

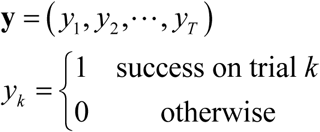

We used the standard LIBSVM toolkit (Chang and Lin 2011) and fed the SVM module with 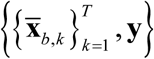, as feature vectors and labels. Training and testing were performed using a 10-fold cross-validation procedure. Trials were divided into 10 equally sized and disjoint sets; in each fold, one of the sets (10% of trials) was used for testing and the remaining 9 sets (90% of the trials) were used for training the classifier. In each fold, the SVM regularization parameter was determined using a grid-search optimization using another (internal) 10-fold partition of the training data. Overall, the 10-fold cross-validation testing process resulted in 10 accuracy rates per time bin *b*, denoted by *a*_*b, f*_, *f* = 1, 2,…,10. We evaluated the mean and STD values across folds as a function of their respective time window *b*,

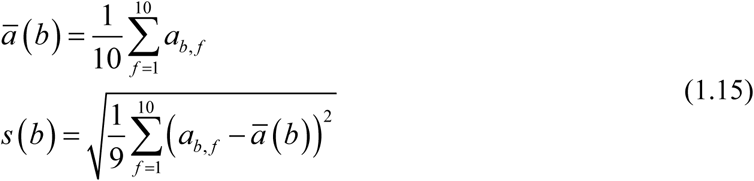

Figure 4F and 4G present the mean and STD values evaluated using eqn. (1.15) as a function of the time bin *b*, where typically the accuracy level starts climbing right after the tone and peaks about 1.5-2 seconds afterwards.

##### Confusion matrices

In the previous section, the effectiveness of the classifier was evaluated by the accuracy rate, which is the amount of trials predicted correctly divided by the overall amount of examples. However, more can be learned about the performance of a classifier from the confusion matrix **M**, where *M* (*i. j*) is the percentage of trials for which examples related to label i were predicted as label j. Therefore, the elements of the main diagonal count the examples correctly classified. In our case, the examples were the trials and the labels were success and failure. Thus, the confusion matrix is a 2-by-2 matrix whose off-diagonal elements consist of the incorrect classification: success classified as failure and vice-versa, as presented in Figure 7B and 7D.

##### Indicative Neurons

In addition to evaluating the separability of success and failure trials based on the neuronal activity of the entire network as a whole, we also aimed to examine whether or not single neurons are able to reliably report success or failure.

We evaluated the average activity of each neuron *i* and time bin *b*, for all trials, as described in eqn. (1.12) and paired them with the associated success and failure labels, 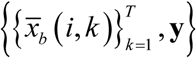, where *i* = 1, 2,…, *N*_*r*_.

We fed the SVM module with 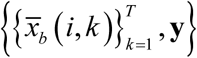 and evaluated the accuracy rate using a 10-fold cross-validation process. In each fold, 90% of the trials were used for training and the remaining 10% were used for testing. The regularization parameter of the SVM was set using an additional 10-fold cross-validation process based on 90% of trials. We obtained the mean and STD values of the accuracy rate, per neuron and at each time window, by averaging over folds,

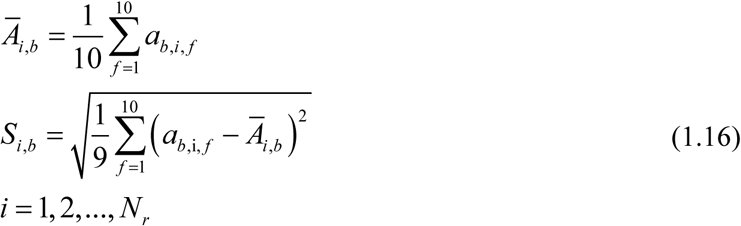

To determine whether or not a neuron is indicative, we compared its respective mean accuracy rate to the prior probability of success and failure, evaluated by

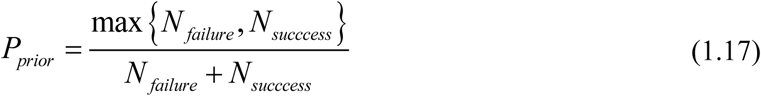

where *N*_*success*_ and *N* _*failure*_ are the number of success and failure trials, respectively. In each time window, we marked a neuron as indicative if its mean accuracy rate was higher than the prior probability, *P*_*prior*_, with a 95% confidence interval, i.e.

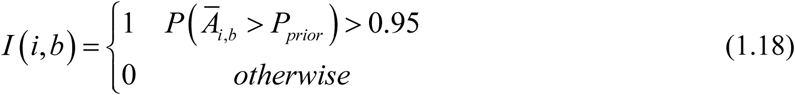

##### Latency histograms

In this section we aimed to temporally correlate between different behavioral events and neuronal activity as measured by calcium imaging. We first extracted time stamps of tone and four behavioral events per trial: the first lift, the first attempt to grab (noted as ‘first grab’), the last attempt to grab (noted as ‘last grab’) and pellet at mouth. We aligned the imaging data per trial (across neurons) according to the time stamps of each of the three events. We obtained the maximal value of neurons with a peak response greater than 3 standard deviations above baseline; the baseline signal was obtained as the mean value of the aligned imaging data taken from 2 seconds before the tone till the tone. We then evaluated the delay between the aligned event (tone, first lift, first grab, last grab or at mouth) and the onset of the calcium transients. Latency histograms depicted in Figure 3A and 3B, present the percentage of neurons activated within a certain delay from the event.

##### GLM

We modeled single neuron calcium transients of both layer 5 PT and layer 2-3 neurons using a Generalized Linear Model (GLM) (Ramkumar et al. 2016, Engelhard et al. 2019). With the GLM we nonlinearly construct predictor signals from the behavioral information of each trial, and then linearly combine them to fit the calcium signal of each neuron.

The predictor signals were of three types:

1. Hand trajectories. Time series of hand trajectories were extracted using DeepLabCut software (Mathis et al. 2018) from videos taken with side and front view cameras. We extracted x and y locations, altogether 4 predictors.
2. Time varying and binary events – tone, lift, grab and at mouth. Movement events were extracted using the modified JAABA software (Kabra et al. 2013).
3. Whole trial binary event – success/failure trial status (outcome).

To model the time course of single neurons calcium signals, we convolved the time varying binary events with a set of 7 degree-of-freedom regression splines. We used a set of short duration (0.5 seconds) and long duration (2 seconds) splines generated using the ‘bSpline’ package in R. We had 16 spline functions in total which resulted in 16 × 4 =64 convolved signals used as predictors. Altogether we had 69 predictors (64 convolved predictors + 4 hand trajectories +1 whole binary success/failure outcome status). We performed our analysis in two time segments: peri-movement – from one second before the tone till 2 seconds after the tone; and post-movement – from 5 seconds after the tone till 8 seconds after the tone. We did not model the intermediate time segment (2-5 seconds after the tone), as in layer 5 PT neurons the calcium transients during this segment were dominated by the residual of the calcium responses evoked by movement events, which occurred during the initial time segment.

On each time segment *τ* we train a linear predictor for the neuronal activity of a neuron *i* based on the corresponding predictors such that:

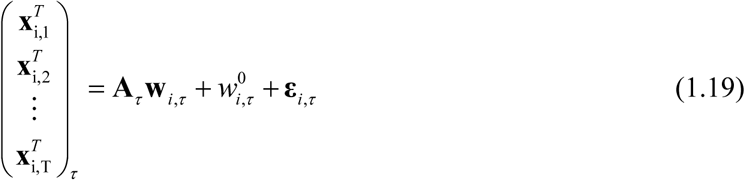

where **x**_*i*,k_ is the temporal neuronal activity of a neuron i on trial k, as defined in (1.1), clipped to the segment *τ*, **A**_*τ*_ consists of the corresponding temporal predictor signals and 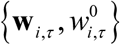 are the evaluated model parameters. All models were trained per neuron and time segment using LASSO (Tibshirani 1996) with 5-fold cross validation.

We first trained a full model based on all 69 predictors and measured 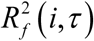, which is the variance of the explained signal normalized by the variance of the neuronal activity.

We analyzed 1008 neurons for layer 5 PT and 1111 for layer 2/3 neurons. For our further analysis we included the subset of neurons having at least 15% of their variance explained by the full model and excluded the rest from further analysis (for layer 5 PT neurons, 552 neurons for the peri-movement segment and none for the post-movement segment; for layer 2/3 neurons, 227 neurons for the peri-movement segment and 78 neurons for the post-movement segment).

In order to quantify the relative contribution of each variable, we grouped the predictors into six categories: tone, lift, grab, at mouth, trajectories and outcome.

For each neuron on each time segment, we trained a set of 6 partial models, each by excluding predictors related to one of the categories. The contribution of the excluded component was evaluated as

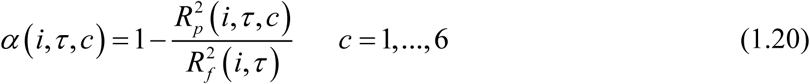

where 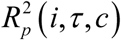 is the variance of the explained signal using the partial model c, c=1,2 …,6.

For some neurons the contribution was negative, indicating poor modeling due to noise or irrelevance of the predictors to the activity and therefore the value was cropped to zero (Engelhard et al. 2019). Figure 3E presents the relative contribution, 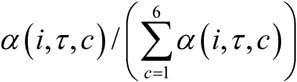 of each component c per time segment *τ*, averaged across neurons.

## DATA AND CODE AVAILABILITY

The code supporting the current study has not been deposited in a public repository but is available from the corresponding author on request.

## References

Adesnik, H., and Naka, A. (2018). Cracking the Function of Layers in the Sensory Cortex. Neuron 100, 1028–1043.

Amador, N., Schlag-Rey, M., and Schlag, J. (2000). Reward-predicting and reward-detecting neuronal activity in the primate supplementary eye field. J Neurophysiol 84, 2166–2170.

Amiez, C., Joseph, J.P., and Procyk, E. (2006). Reward encoding in the monkey anterior cingulate cortex. Cereb Cortex 16, 1040–1055.

Anderson, C.T., Sheets, P.L., Kiritani, T., and Shepherd, G.M. (2010). Sublayer-specific microcircuits of corticospinal and corticostriatal neurons in motor cortex. Nat Neurosci 13, 739–744.

Bosch-Bouju, C., Hyland, B.I., and Parr-Brownlie, L.C. (2013). Motor thalamus integration of cortical, cerebellar and basal ganglia information: implications for normal and parkinsonian conditions. Front Comput Neurosci 7, 163.

Chen, T.W., Li, N., Daie, K., and Svoboda, K. (2017). A Map of Anticipatory Activity in Mouse Motor Cortex. Neuron 94, 866–879 e864.

Churchland, M.M., Cunningham, J.P., Kaufman, M.T., Foster, J.D., Nuyujukian, P., Ryu, S.I., and Shenoy, K.V. (2012). Neural population dynamics during reaching. Nature 487, 51–56.

Cichon, J., and Gan, W.B. (2015). Branch-specific dendritic Ca(2+) spikes cause persistent synaptic plasticity. Nature 520, 180–185.

Cunningham, J.P., and Yu, B.M. (2014). Dimensionality reduction for large-scale neural recordings. Nat Neurosci 17, 1500–1509.

Ebbesen, C.L., and Brecht, M. (2017). Motor cortex - to act or not to act? Nat Rev Neurosci 18, 694–705.

Engelhard, B., Finkelstein, J., Cox, J., Fleming, W., Jang, H.J., Ornelas, S., Koay, S.A., Thiberge, S.Y., Daw, N.D., Tank, D.W., et al. (2019). Specialized coding of sensory, motor and cognitive variables in VTA dopamine neurons. Nature 570, 509–513.

Even-Chen, N., Stavisky, S.D., Kao, J.C., Ryu, S.I., and Shenoy, K.V. (2017). Augmenting intracortical brain-machine interface with neurally driven error detectors. J Neural Eng 14, 066007.

Friston, K. (2011). What is optimal about motor control? Neuron 72, 488–498.

Fu, M., Yu, X., Lu, J., and Zuo, Y. (2012). Repetitive motor learning induces coordinated formation of clustered dendritic spines in vivo. Nature 483, 92–95.

Galea, J.M., Mallia, E., Rothwell, J., and Diedrichsen, J. (2015). The dissociable effects of punishment and reward on motor learning. Nat Neurosci 18, 597–602.

Georgopoulos, A.P., and Carpenter, A.F. (2015). Coding of movements in the motor cortex. Curr Opin Neurobiol 33, 34–39.

Glascher, J., Daw, N., Dayan, P., and O’Doherty, J.P. (2010). States versus rewards: dissociable neural prediction error signals underlying model-based and model-free reinforcement learning. Neuron 66, 585–595.

Guo, J.Z., Graves, A.R., Guo, W.W., Zheng, J., Lee, A., Rodriguez-Gonzalez, J., Li, N., Macklin, J.J., Phillips, J.W., Mensh, B.D., et al. (2015). Cortex commands the performance of skilled movement. eLife 4, e10774.

Harris, K.D., and Shepherd, G.M. (2015). The neocortical circuit: themes and variations. Nat Neurosci 18, 170–181.

Hayashi-Takagi, A., Yagishita, S., Nakamura, M., Shirai, F., Wu, Y.I., Loshbaugh, A.L., Kuhlman, B., Hahn, K.M., and Kasai, H. (2015). Labelling and optical erasure of synaptic memory traces in the motor cortex. Nature 525, 333–338.

Heffley, W., Song, E.Y., Xu, Z., Taylor, B.N., Hughes, M.A., McKinney, A., Joshua, M., and Hull, C. (2018). Coordinated cerebellar climbing fiber activity signals learned sensorimotor predictions. Nat Neurosci 21, 1431–1441.

Heindorf, M., Arber, S., and Keller, G.B. (2018). Mouse Motor Cortex Coordinates the Behavioral Response to Unpredicted Sensory Feedback. Neuron 99, 1040–1054 e1045.

Hooks, B.M., Hires, S.A., Zhang, Y.X., Huber, D., Petreanu, L., Svoboda, K., and Shepherd, G.M. (2011). Laminar analysis of excitatory local circuits in vibrissal motor and sensory cortical areas. PLoS Biol 9, e1000572.

Hooks, B.M., Mao, T., Gutnisky, D.A., Yamawaki, N., Svoboda, K., and Shepherd, G.M. (2013). Organization of cortical and thalamic input to pyramidal neurons in mouse motor cortex. J Neurosci 33, 748–760.

Hosp, J.A., Pekanovic, A., Rioult-Pedotti, M.S., and Luft, A.R. (2011). Dopaminergic projections from midbrain to primary motor cortex mediate motor skill learning. J Neurosci 31, 2481–2487.

Huang, C.C., Sugino, K., Shima, Y., Guo, C., Bai, S., Mensh, B.D., Nelson, S.B., and Hantman, A.W. (2013). Convergence of pontine and proprioceptive streams onto multimodal cerebellar granule cells. eLife 2, e00400.

Huber, D., Gutnisky, D.A., Peron, S., O’Connor, D.H., Wiegert, J.S., Tian, L., Oertner, T.G., Looger, L.L., and Svoboda, K. (2012). Multiple dynamic representations in the motor cortex during sensorimotor learning. Nature 484, 473–478.

Inoue, M., Uchimura, M., and Kitazawa, S. (2016). Error Signals in Motor Cortices Drive Adaptation in Reaching. Neuron 90, 1114–1126.

Isomura, Y., Harukuni, R., Takekawa, T., Aizawa, H., and Fukai, T. (2009). Microcircuitry coordination of cortical motor information in self-initiation of voluntary movements. Nat Neurosci 12, 1586–1593.

Isomura, Y., Takekawa, T., Harukuni, R., Handa, T., Aizawa, H., Takada, M., and Fukai, T. (2013). Reward-modulated motor information in identified striatum neurons. J Neurosci 33, 10209–10220.

Izawa, J., and Shadmehr, R. (2011). Learning from sensory and reward prediction errors during motor adaptation. PLoS Comput Biol 7, e1002012.

Kabra, M., Robie, A.A., Rivera-Alba, M., Branson, S., and Branson, K. (2013). JAABA: interactive machine learning for automatic annotation of animal behavior. Nat Methods 10, 64–67.

Keller, G.B., Bonhoeffer, T., and Hubener, M. (2012). Sensorimotor mismatch signals in primary visual cortex of the behaving mouse. Neuron 74, 809–815.

Keller, G.B., and Mrsic-Flogel, T.D. (2018). Predictive Processing: A Canonical Cortical Computation. Neuron 100, 424–435.

Kojima, S., Kao, M.H., Doupe, A.J., and Brainard, M.S. (2018). The Avian Basal Ganglia Are a Source of Rapid Behavioral Variation That Enables Vocal Motor Exploration. J Neurosci 38, 9635–9647.

Komiyama, T., Sato, T.R., O’Connor, D.H., Zhang, Y.X., Huber, D., Hooks, B.M., Gabitto, M., and Svoboda, K. (2010). Learning-related fine-scale specificity imaged in motor cortex circuits of behaving mice. Nature 464, 1182–1186.

Kording, K.P., and Wolpert, D.M. (2006). Bayesian decision theory in sensorimotor control. Trends Cogn Sci 10, 319–326.

Kostadinov, D., Beau, M., Pozo, M.B., and Hausser, M. (2019). Predictive and reactive reward signals conveyed by climbing fiber inputs to cerebellar Purkinje cells. Nat Neurosci 22, 950–962.

Krigolson, O.E., and Holroyd, C.B. (2007). Predictive information and error processing: the role of medial-frontal cortex during motor control. Psychophysiology 44, 586–595.

Kuramoto, E., Furuta, T., Nakamura, K.C., Unzai, T., Hioki, H., and Kaneko, T. (2009). Two types of thalamocortical projections from the motor thalamic nuclei of the rat: a single neurontracing study using viral vectors. Cereb Cortex 19, 2065–2077.

Lacefield, C.O., Pnevmatikakis, E.A., Paninski, L., and Bruno, R.M. (2019). Reinforcement Learning Recruits Somata and Apical Dendrites across Layers of Primary Sensory Cortex. Cell Rep 26, 2000–2008 e2002.

Lafon, S., Keller, Y., and Coifman, R.R. (2006). Data fusion and multicue data matching by diffusion maps. IEEE Trans Pattern Anal Mach Intell 28, 1784–1797.

Laubach, M., Wessberg, J., and Nicolelis, M.A. (2000). Cortical ensemble activity increasingly predicts behaviour outcomes during learning of a motor task. Nature 405, 567–571.

Lemon, R.N. (2008). Descending pathways in motor control. Annu Rev Neurosci 31, 195–218.

Li, Q., Ko, H., Qian, Z.M., Yan, L.Y.C., Chan, D.C.W., Arbuthnott, G., Ke, Y., and Yung, W.H. (2017). Refinement of learned skilled movement representation in motor cortex deep output layer. Nat Commun 8, 15834.

Mahn, M., Gibor, L., Patil, P., Cohen-Kashi Malina, K., Oring, S., Printz, Y., Levy, R., Lampl, I., and Yizhar, O. (2018). High-efficiency optogenetic silencing with soma-targeted anion-conducting channelrhodopsins. Nat Commun 9, 4125.

Makino, H., Hwang, E.J., Hedrick, N.G., and Komiyama, T. (2016). Circuit Mechanisms of Sensorimotor Learning. Neuron 92, 705–721.

Mao, T., Kusefoglu, D., Hooks, B.M., Huber, D., Petreanu, L., and Svoboda, K. (2011). Long-range neuronal circuits underlying the interaction between sensory and motor cortex. Neuron 72, 111–123.

Masamizu, Y., Tanaka, Y.R., Tanaka, Y.H., Hira, R., Ohkubo, F., Kitamura, K., Isomura, Y., Okada, T., and Matsuzaki, M. (2014). Two distinct layer-specific dynamics of cortical ensembles during learning of a motor task. Nat Neurosci 17, 987–994.

Mathis, A., Mamidanna, P., Cury, K.M., Abe, T., Murthy, V.N., Mathis, M.W., and Bethge, M. (2018). DeepLabCut: markerless pose estimation of user-defined body parts with deep learning. Nat Neurosci 21, 1281–1289.

Mathis, M.W., Mathis, A., and Uchida, N. (2017). Somatosensory Cortex Plays an Essential Role in Forelimb Motor Adaptation in Mice. Neuron 93, 1493–1503 e1496.

Matsumoto, M., Matsumoto, K., Abe, H., and Tanaka, K. (2007). Medial prefrontal cell activity signaling prediction errors of action values. Nat Neurosci 10, 647–656.

Molina-Luna, K., Pekanovic, A., Rohrich, S., Hertler, B., Schubring-Giese, M., Rioult-Pedotti, M.S., and Luft, A.R. (2009). Dopamine in motor cortex is necessary for skill learning and synaptic plasticity. PLoS One 4, e7082.

Nikooyan, A.A., and Ahmed, A.A. (2015). Reward feedback accelerates motor learning. J Neurophysiol 113, 633–646.

Osborne, J.E., and Dudman, J.T. (2014). RIVETS: a mechanical system for in vivo and in vitro electrophysiology and imaging. PLoS One 9, e89007.

Papale, A.E., and Hooks, B.M. (2018). Circuit changes in motor cortex during motor skill learning. Neuroscience 368, 283–297.

Peters, A.J., Chen, S.X., and Komiyama, T. (2014). Emergence of reproducible spatiotemporal activity during motor learning. Nature 510, 263–267.

Peters, A.J., Lee, J., Hedrick, N.G., O’Neil, K., and Komiyama, T. (2017). Reorganization of corticospinal output during motor learning. Nat Neurosci 20, 1133–1141.

Petrof, I., Viaene, A.N., and Sherman, S.M. (2015). Properties of the primary somatosensory cortex projection to the primary motor cortex in the mouse. J Neurophysiol 113, 2400–2407.

Ramkumar, P., Dekleva, B., Cooler, S., Miller, L., and Kording, K. (2016). Premotor and Motor Cortices Encode Reward. PLoS One 11, e0160851.

Raymond, J.L., and Medina, J.F. (2018). Computational Principles of Supervised Learning in the Cerebellum. Annu Rev Neurosci 41, 233–253.

Roesch, M.R., and Olson, C.R. (2004). Neuronal activity related to reward value and motivation in primate frontal cortex. Science 304, 307–310.

Roesch, M.R., and Olson, C.R. (2005). Neuronal activity in primate orbitofrontal cortex reflects the value of time. J Neurophysiol 94, 2457–2471.

Sajad, A., Godlove, D.C., and Schall, J.D. (2019). Cortical microcircuitry of performance monitoring. Nat Neurosci 22, 265–274.

Schall, J.D., Stuphorn, V., and Brown, J.W. (2002). Monitoring and control of action by the frontal lobes. Neuron 36, 309–322.

Schultz, W. (2000). Multiple reward signals in the brain. Nat Rev Neurosci 1, 199–207.

Shadmehr, R., and Krakauer, J.W. (2008). A computational neuroanatomy for motor control. Exp Brain Res 185, 359–381.

Shepherd, G.M. (2013). Corticostriatal connectivity and its role in disease. Nat Rev Neurosci 14, 278–291.

Shmuelof, L., and Krakauer, J.W. (2011). Are we ready for a natural history of motor learning? Neuron 72, 469–476.

Stuphorn, V., Taylor, T.L., and Schall, J.D. (2000). Performance monitoring by the supplementary eye field. Nature 408, 857–860.

Teichert, T., Yu, D., and Ferrera, V.P. (2014). Performance monitoring in monkey frontal eye field. J Neurosci 34, 1657–1671.

Tervo, D.G., Hwang, B.Y., Viswanathan, S., Gaj, T., Lavzin, M., Ritola, K.D., Lindo, S., Michael, S., Kuleshova, E., Ojala, D., et al. (2016). A Designer AAV Variant Permits Efficient Retrograde Access to Projection Neurons. Neuron 92, 372–382.

Tseng, Y.W., Diedrichsen, J., Krakauer, J.W., Shadmehr, R., and Bastian, A.J. (2007). Sensory prediction errors drive cerebellum-dependent adaptation of reaching. J Neurophysiol 98, 54–62.

Tsubo, Y., Isomura, Y., and Fukai, T. (2013). Neural dynamics and information representation in microcircuits of motor cortex. Front Neural Circuits 7, 85.

Uehara, S., Mawase, F., Therrien, A.S., Cherry-Allen, K.M., and Celnik, P. (2019). Interactions between motor exploration and reinforcement learning. J Neurophysiol 122, 797–808.

Wagner, M.J., Kim, T.H., Savall, J., Schnitzer, M.J., and Luo, L. (2017). Cerebellar granule cells encode the expectation of reward. Nature 544, 96–100.

Wallis, J.D., and Kennerley, S.W. (2010). Heterogeneous reward signals in prefrontal cortex. Curr Opin Neurobiol 20, 191–198.

Watabe-Uchida, M., Eshel, N., and Uchida, N. (2017). Neural Circuitry of Reward Prediction Error. Annu Rev Neurosci 40, 373–394.

Weiler, N., Wood, L., Yu, J., Solla, S.A., and Shepherd, G.M. (2008). Top-down laminar organization of the excitatory network in motor cortex. Nat Neurosci 11, 360–366.

Whishaw, I.Q. (2000). Loss of the innate cortical engram for action patterns used in skilled reaching and the development of behavioral compensation following motor cortex lesions in the rat. Neuropharmacology 39, 788–805.

Whishaw, I.Q., O’Connor, W.T., and Dunnett, S.B. (1986). The contributions of motor cortex, nigrostriatal dopamine and caudate-putamen to skilled forelimb use in the rat. Brain 109 (Pt 5), 805–843.

Wickens, J.R., Reynolds, J.N., and Hyland, B.I. (2003). Neural mechanisms of reward-related motor learning. Curr Opin Neurobiol 13, 685–690.

Wolpert, D.M., Diedrichsen, J., and Flanagan, J.R. (2011). Principles of sensorimotor learning. Nat Rev Neurosci 12, 739–751.

Wolpert, D.M., and Miall, R.C. (1996). Forward Models for Physiological Motor Control. Neural Netw 9, 1265–1279.

Zeng, H., and Sanes, J.R. (2017). Neuronal cell-type classification: challenges, opportunities and the path forward. Nat Rev Neurosci 18, 530–546.

## References

1. D. G. Tervo et al., A Designer AAV Variant Permits Efficient Retrograde Access to Projection Neurons. Neuron 92, 372–382 (2016).

2. G. Silasi, J. D. Boyd, J. Ledue, T. H. Murphy, Improved methods for chronic light-based motor mapping in mice: automated movement tracking with accelerometers, and chronic EEG recording in a bilateral thin-skull preparation. Front Neural Circuits 7, 123 (2013).

3. J. E. Osborne, J. T. Dudman, RIVETS: a mechanical system for in vivo and in vitro electrophysiology and imaging. PLoS One 9, e89007 (2014).

4. Mahn, Mathias, Lihi Gibor, Pritish Patil, Katayun Cohen-Kashi Malina, Shir Oring, Yoav Printz, Rivka Levy, Ilan Lampl, and Ofer Yizhar. High-efficiency optogenetic silencing with soma-targeted anion-conducting channelrhodopsins. Nature communications 9 (2018).

5. M. Kabra, A. A. Robie, M. Rivera-Alba, S. Branson, K. Branson, JAABA: interactive machine learning for automatic annotation of animal behavior. Nat Methods 10, 64–67 (2013).

6. F. J. R. Kowalczyk A. M., Phase retrival for a complex-valued object by using a low-resolution image. J. Opt. Soc. Am., 450–458 (1990).

7. C.-C. Chang and C.-J. Lin. LIBSVM: A library for support vector machines. ACM Transactions on Intelligent Systems and Technology, 2:1{27, 2011. Software available at http://www.csie.ntu.edu.tw/~cjlin/libsvm.

8. Mishne, G., Talmon, R., Meir, R., Schiller, J., Lavzin, M., Dubin, U. and Coifman, R.R., 2016. Hierarchical coupled-geometry analysis for neuronal structure and activity pattern discovery. IEEE Journal of Selected Topics in Signal Processing, 10(7), pp.1238–1253.

9. Belkin M, Niyogi P. Laplacian eigenmaps for dimensionality reduction and data representation. Neural computation. 2003 Jun 1;15(6):1373–96.

10. Donoho DL, Grimes C. Hessian eigenmaps: Locally linear embedding techniques for high-dimensional data. Proceedings of the National Academy of Sciences. 2003 May 13;100(10):5591–6.

11. Coifman RR, Lafon S, Lee AB, Maggioni M, Nadler B, Warner F, Zucker SW. Geometric diffusions as a tool for harmonic analysis and structure definition of data: Diffusion maps. Proceedings of the national academy of sciences. 2005 May 24;102(21):7426–31.

12. Coifman, R.R. and Lafon, S., 2006. Diffusion maps. Applied and computational harmonic analysis, 21(1), pp.5–30.

13. Mishne, G., Talmon, R., Cohen, I., Coifman, R.R. and Kluger, Y., 2017. Data-driven tree transforms and metrics. arXiv preprint 1708.05768.

14. Shemesh, A., Talmon, R., Karp, O., Amir, I., Bar, M. and Grobman, Y.J., 2017. Affective response to architecture–investigating human reaction to spaces with different geometry. Architectural Science Review, 60(2), pp.116–125.

15. Yair, O. and Talmon, R., 2017. Local canonical correlation analysis for nonlinear common variables discovery. IEEE Transactions on Signal Processing, 65(5), pp.1101–1115.

16. Shnitzer, T., Talmon, R. and Slotine, J.J., 2016. Manifold learning with contracting observers for data-driven time-series analysis. arXiv preprint 1604.04492.

17. Sulam, J., Romano, Y. and Talmon, R., 2017. Dynamical system classification with diffusion embedding for ECG-based person identification. Signal Processing, 130, pp.403–411.

18. Joachims, T., 2006, August. Training linear SVMs in linear time. In Proceedings of the 12th ACM SIGKDD international conference on Knowledge discovery and data mining (pp. 217–226). ACM.

19. Tibshirani, R. Regression shrinkage and selection via the lasso. Journal of the Royal Statistical Society: Series B (Methodological) 58.1 (1996): 267–288.

20. Ramkumar, Pavan, et al. Premotor and motor cortices encode reward. PloS one 11.8 (2016): e0160851.

21. Engelhard, Ben, Joel Finkelstein, Julia Cox, Weston Fleming, Hee Jae Jang, Sharon Ornelas, Sue Ann Koay et al. Specialized coding of sensory, motor and cognitive variables in VTA dopamine neurons. Nature (2019): 1.

22. Mathis, Alexander, et al. DeepLabCut: markerless pose estimation of user-defined body parts with deep learning. Nature Publishing Group, 2018.

